# *In silico* Exploration Natural Compounds for the Discovery of Novel DNMT3A Inhibitors as Potential Therapeutic Agents for Acute Myeloid Leukemia

**DOI:** 10.1101/2024.08.28.610083

**Authors:** Uddalak Das, Akshay Uttarkar, Jitendra Kumar, Vidya Niranjan

## Abstract

Aberrant DNA methylation, a hallmark of acute myeloid leukemia (AML), is catalyzed by DNA methyltransferase 3A (DNMT3A). Approximately 20-30% of AML patients harbor DNMT3A mutations, leading to disrupted DNA methylation patterns and leukemogenesis. To identify potential therapeutic interventions, this study employed computational drug discovery. A pharmacophore model was constructed and utilized to screen a natural product database, yielding a set of promising compounds. Subsequent molecular docking, MM-GBSA calculations, and ADMET profiling identified two compounds, CNP0375130 and CNP0256178, as potential DNMT3A inhibitors. These compounds exhibited favorable binding affinities and demonstrated desirable drug-like properties. Molecular dynamics simulations confirmed stable protein-ligand interactions. These findings suggest that CNP0375130 and CNP0256178 may serve as promising lead compounds for the development of novel anti-leukemic therapies targeting DNMT3A, and contribute to the ongoing efforts to develop targeted therapies for leukemia.

## 1. INTRODUTION

Blood cancer, arising from blood-forming cells, bone marrow, and the lymphatic system, encompasses various types such as leukemia, lymphoma, and myeloma, with leukemia ranking 10^th^ deadliest among common cancers (Sung *et al*., 2021). Epigenetic dysregulation, involving heritable changes in gene expression without altering DNA sequence, plays a pivotal role in cancer, including leukemia (Sharma *et al*., 2010). DNA methylation, a crucial epigenetic mechanism, regulates genome expression and homeostasis. Dysregulation, leading to aberrant hypermethylation or hypomethylation, disrupts normal cellular functions and is implicated in cancer initiation and progression (A. Klupczyńska, 2023). Central to this process are DNA methyltransferases (DNMTs), including DNMT1, DNMT3A, and DNMT3B. DNMT1 maintains methylation patterns during DNA replication, while DNMT3A and DNMT3B catalyze de novo methylation (Jin & Robertson, 2013).

DNMTs are enzymes that catalyze DNA methylation by transferring a methyl group from S-Adenosyl-L-Methionine (SAM) to the cytosine ring of DNA, producing S-Adenosyl-L-Homocysteine (SAH) (Jin & Robertson, 2013). Aberrant DNA hypermethylation plays a crucial role in leukemogenesis. Nordlund *et al*. (2013) demonstrated widespread hypermethylation across CpG sites in children with ALL and at relapse stages, implicating DNA methylation in disease progression (Nordlund *et al*., 2013). Similarly, hypermethylation has been associated with the progression from Myelodysplastic Syndromes (MDS) to Acute Myeloid Leukemia (AML) and influences the pathogenesis of other leukemia types such as Chronic Myeloid Leukemia (CML) and ALL (Kalinkova *et al*., 2022). DNMT3A mutations are prevalent in approximately 20% of AML cases and are associated with altered DNA methylation patterns (Park *et al*., 2020). DNMT3A mutations, predominantly heterozygous, often cluster at the methyltransferase domain, disrupting normal methylation activity (L. Yang *et al*., 2015). These mutations have been implicated as founder mutations in AML and are frequently observed alongside mutations in genes like NPM1 and FLT3, influencing leukemic transcription factors and clinical outcomes (Oñate *et al*., 2022).

*In silico* drug discovery has emerged as a time-efficient and cost-effective approach for screening therapeutic compounds. This method combines various computational techniques such as molecular docking and molecular dynamics (MD) simulations to predict the binding affinity and stability of protein-ligand complexes (Chang *et al*., 2022). Molecular docking helps in the initial screening of multiple compounds based on their binding capacity to the target protein, while MD simulations provide insights into the stability of these complexes in a virtual environment that mimics the human body (Meng *et al*., 2011).

In this study, we employed a combination of ligand-based pharmacophore modeling, molecular docking, and MMGBSA to screen and validate potential DNMT3A inhibitors from the COCONUT database. The study also involved ADMET profiling and molecular dynamics simulations to assess the stability and drug-likeness of the identified compounds. By targeting DNMT3A, we aim to identify novel inhibitors that could serve as potential therapeutic agents for the treatment of leukemia. The insights gained from our research will contribute to the development of more potent DNMT3A inhibitors, advancing the field of epigenetic therapy for leukemia.

## 2. MATERIALS AND METHODS

### 2.1. Dataset generation and screening

The COCONUT (version 2022, Collection of Open Natural ProdUcTs Online) database was utilized as the screening source, containing approximately 407,270 molecules (Sorokina *et al*., 2021). The molecules were then filtered using QikProp module in “Structure Filtering” option Schrödinger 2023-1, with zero violation of Lipinski’s Rule of 5 generating an output of 276409 molecules. The filtered structures where then prepared using the LigPrep Module with the OPLS4 force field (Lu *et al*., 2021). The possible states of molecules were generated using Epik at pH 7.0 ± 2.0, retained specified chirality, followed by tautomer generation and the generation of up to 16 low-energy conformations per ligand. The total number of generated structures were around 3.57 million.

### 2.2. Pharmacophore modelling/screening

#### 2.2.1. Pharmacophore hypothesis generation

A pharmacophore comprises distinct characteristics like steric properties, hydrogen bond donor (HBD), hydrogen bond acceptor (HBA), and electronic chemical attributes. These features indicate how a compound functions uniquely within the active biological site (Choudhury & Narahari Sastry, 2019). The development of ligand-based pharmacophore design relied on identified actives with confirmed pharmacological effects targeting the chosen receptor.

To generate the pharmacophore model, an initial subset comprising the first 100 compounds linked to the DNMT3A was extracted from the binding database, sorted based on their respective IC_50_ values (Ganji *et al*., 2023). This curated dataset was the foundation for subsequent pharmacophore model construction and screening endeavors. Within the LigPrep module, the OPLS4 force field was engaged (Lu *et al*., 2021), concomitant with default parameters for ionization. Furthermore, routine procedures encompassing desalination, tautomerization, and computational adjustments were implemented per software defaults. It helps to prepare high-quality 3D structures for drug-like molecules.

In the Develop Pharmacophore Model module, the hypothesis match was set to 25% given the high diversity of ligands obtained from BindingDB and the number of features in the hypothesis was kept from 4 to 7 with the preferred number to 5. The ranking and scoring of the hypothesis were set to the default “Phase Hypo Score” (Yu *et al*., 2021). The Generate Conformer and Minimize Output conformer options were activated, with the target number of conformers set to 50 (Cole *et al*., 2018).

#### 2.2.2. Validation of the pharmacophore model

The developed pharmacophore model was verified to check its ability to predict the activity of new compounds effectively. This procedure is a prerequisite before the pharmacophore model can be employed for virtual screening (Kaserer *et al*., 2015). The parameters used for evaluating the efficiency of the developed pharmacophore model are enrichment factor (EF), receiver operating characteristic (ROC) curves, Boltzmann-enhanced discrimination of ROC (BEDROC) (Truchon & Bayly, 2007), and robust initial enhancement (RIE) (Truchon & Bayly, 2007).

A decoy set was created using the Generate DUDE Decoys program, which is found at http://dude.docking.org/generate (Mysinger *et al*., 2012). For converting the output into 3D structures, Open Babel v2.4.1 was used (O’Boyle *et al*., 2011). Ligand preparation was done following the default settings and the protocols of LigPrep. The OPLS4 force field has been employed in the minimization procedure (Lu *et al*., 2021).

#### 2.2.3. Screening using Pharmacophore Model

The prepared ligands were screened using the best generated pharmacophore model, AAHR-1 using the “Phase Module”. Prefer partial matches of more features were activated. All the pharmacophore properties (4 of 4) were selected for screening. The pharmacophore model screened ∼3.57 million structures to ∼220K structures.

### 2.3. Molecular Interactions

#### 2.3.1. Protein preparation

The X-ray crystallographic structure of the DNMT3A target protein [PDB ID: 5YX2(A)], found at https://www.rcsb.org, with a resolution of 2.65Å, was used in the study (Zhang *et al*., 2018). The structure analysis was carried out using the PDBsum web server (https://www.ebi.ac.uk/thornton-srv/databases/pdbsum). PDBsum online server was also used to check the validation of the DNMT3A with the Ramachandran plot (Laskowski *et al*., 2018).

In the Schrödinger Maestro protein preparation wizard, the protein was pre-processed with the PROPKA module for an optimization of H-bonds (Gokcan & Isayev, 2022), followed by minimization of structures towards convergence of heavy atoms at RMSD 0.3Å using OPLS4 force field and removal of water molecules more than 5Å away from ligands afterward (Lu *et al*., 2021).

#### 2.3.2. Receptor grid generation

The receptor grid was generated keeping the hydrophobic region and also the region where S-adenosyl-L-Homocysteine (SAH) was attached to the complex, at the centroid of the grid. The coordinates of the receptor grid were X=60.49, Y=32.38, Z=-21.38, with ligand size upto 18Å

#### 2.3.3. Molecular docking

Docking was limited to ligands with 100 rotatable bonds and fewer than 500 atoms. Van der Waals radii scaling factor was set to 0.80, with a partial charge cutoff of 0.15 (Pavlin *et al*., 2019). Sample nitrogen inversions and sample ring conformations were activated, and the ligand sampling was set to flexible. All predicted functional groups had bias sampling of torsions enabled. The module was configured to promote intramolecular hydrogen bonds and improve conjugated pi groups’ planarity. Extra precision (XP) docking was done using the Glide module of Schrödinger.

Docking was done through a series of hierarchical filters i.e. HTVS mode (high-throughput virtual screening) for efficiently screening million compound libraries, to the SP mode (standard precision) for reliably docking tens to hundreds of thousands of ligands with high accuracy, to the XP mode (extra precision) where further elimination of false positives is accomplished by more extensive sampling and advanced scoring, resulting in even higher enrichment. Each step proceeded with the top 10% from the previous one (Bagchi *et al*., 2017). The HTVS filtered the library to 89,386, SP filtered to 8,876 and, XP filtered to around 891 structures respectively.

#### 2.3.4. MM-GBSA Screening

The prime module was utilized to predict the energy parameters obtained from the MM-GBSA (Molecular Mechanics-Generalized Born Surface Area) simulation. It aimed to predict the amounts of the stabilization energy coming from the potential interaction between the selected ligands and the target receptor 5YX2(A) (Genheden & Ryde, 2015). The VGSB solvation model was used and the force field was set as OPLS4 (Lu *et al*., 2021).

The top 10% of the ligands generated through XP docking were analysed using MM-GBSA. The top 2 compounds – CNP0375130 (−76.41 kcal/mol) and CNP0256178 (−75.02 kcal/mol) were used for further ADME-Tox and MD Simulation studies.

### 2.4. *In silico* ADME/T and toxicity analysis

QikProp Module of Schrödinger, SwissADME (Daina *et al*., 2017), ProTox-3.0 (Banerjee *et al*., 2024), ADMETlab 13.0 (Xiong *et al*., 2021), and pkCSM (Pires *et al*., 2015) server were used in the analysis of pharmacokinetic properties to assess the detailed ADMET properties of the two best-ranked compounds based on lowest binding evergy from MM-GBSA screening. SwissADME, pkCSM and ProTox-3.0 are free web tools for predicting pharmacokinetics, drug-likeness and medicinal chemistry friendliness of small molecules.

### 2.5. Molecular Dynamics (MD) simulation studies

Desmond package was used to carry out the molecular dynamics simulations for the 5YX2(A)-CNP0375130 and 5YX2(A)-CNP0256178 complexes. Each system was placed individually in an orthorhombic water box of 10 Å using the TIP3P water model (Mark & Nilsson, 2001). The ligand-protein complexes were modelled by the OPLS4 force field (Lu *et al*., 2021). Counter ions (Na+) were introduced in the ligand-protein complex structures to neutralize the total charge of the systems undergoing MD simulation. Furthermore, the energies of the systems were minimized to a minimum level using 2000 steps before initiating the MD simulation along an NPT lattice trajectory (Al-Jumaili *et al*., 2023).

The RMSD primarily suggests the stability of the ligand interaction, while RMSF describes the fluctuation and flexibility of the residues within the protein, particularly within the active site that is crucial for drug discovery 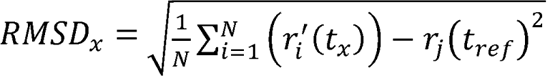. RMSD is calculated by the discovery. RMSF values also help in defining protein structures as they provide information about local conformational changes within the protein chains. (Bharadwaj, Bhargava, & Ball, 2021). The RMSF values are in units of Å and is calculated by the following equation 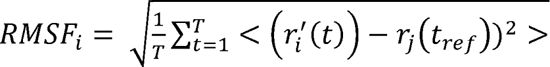 (Ghahremanian *et al*., 2022)

### 2.6. Principal Component Analysis (PCA) and Dynamical Cross-Correlation Matrix (DCCM) Analysis

The trajectory file generated after the MD simulation was first extracted as a .cms file. This .cms file was then converted to .dcd format with the help of VMD 1.9.3 software (Humphrey *et al*., 1996). The trajectory .dcd file was subsequently uploaded to the MDM-TASK web server to perform PCA and DCCM analysis (Amamuddy *et al*., 2021). PCA was conducted to identify the major conformational changes and essential dynamics of the protein-ligand complex, while DCCM analysis was used to examine the correlated motions of residues over the simulation period.

### 2.7. MM-GBSA analysis

The MM-GBSA-based binding free energy calculations were done on the 100ns long MDS trajectories. For the selection of protein-ligand complexes, the binding energies calculated by this approach are more efficient than the glide score values. The main energy elements like H-bond interaction energy (ΔG_Bind_Hbond_), electrostatic solvation free energy (ΔG_Bind_Solv_), Coulomb or electrostatics interaction energy (ΔG_Bind_Coul_), lipophilic interaction energy (ΔG_Bind_Lipo_), and van der Waals interaction energy (ΔG_Bind_vdW_) altogether were considered to the calculation of MM-GBSA based relative binding affinity.

## 3. RESULTS

### 3.1. The DNMT3A Protein

Analysis through PDBsum web server identified 14 α-helices, 3 β-hairpins, 1 β-bulges, 26 β-turns as well as 14 helix-helix interactions. Furthermore, the Ramachandran plot was also used to validate DNMT3A, and it indicated that 90.3% of the residues were in preferred regions, 9.3% were in additional residue regions, 0.1% in generous regions, and 0.2% in disallowed regions with a total G-Factor of 0.33. (Ramachandran *et. al.,* 1963) (see Figure 2)

**Figure 1.**
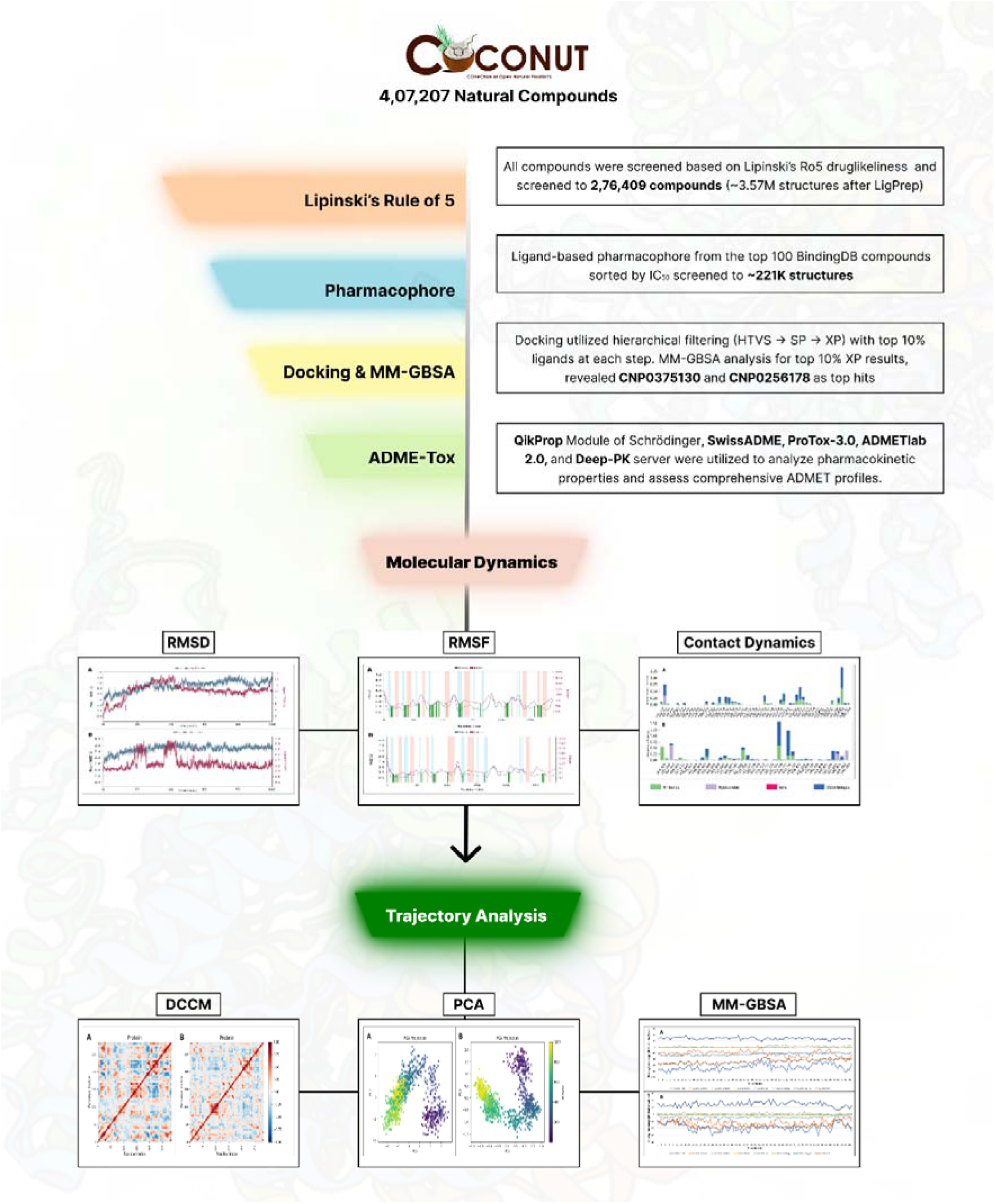
The schematic diagram of the complete dataset screening steps used in this study (Graphical Abstract)

**Figure 2.**
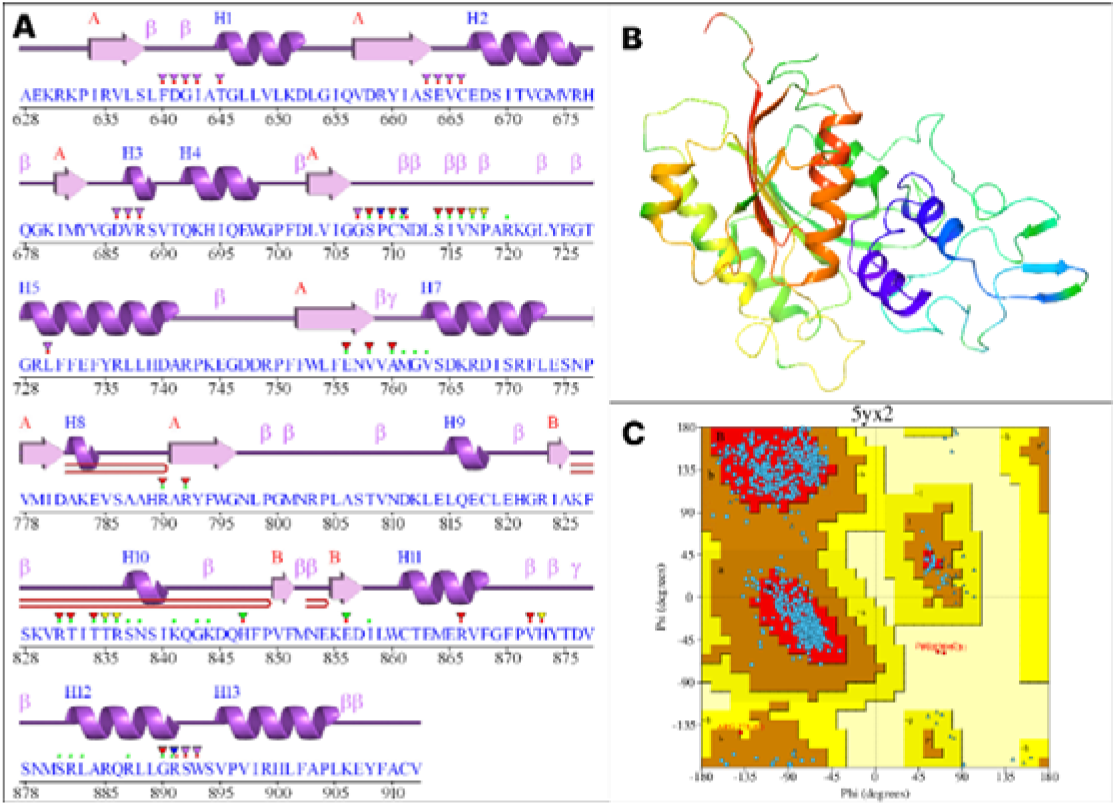
A: The secondary structure of the amino acid sequence of protein 5YX2(A) (generated by PDBsum); B: The 3D structure of the DNMT3A (Chain A) protein; C: Ramachandran plot of the protein. The colour red indicates Iow-energy regions, yellow - allowed regions, pale yellow - generously allowed regions, and white - disallowed regions.

**Figure 3.**
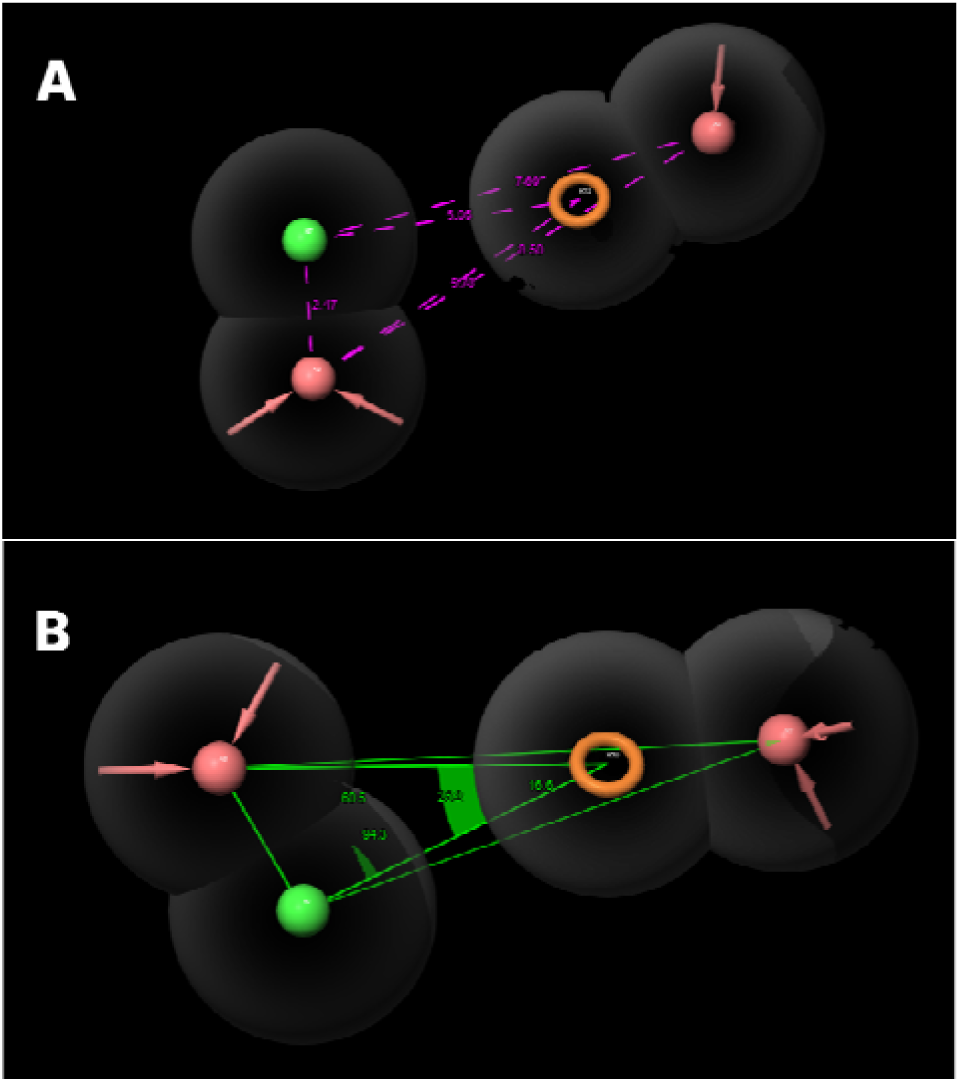
The distance (A) and angles (B) between pharmacophore features of the best pharmacophore model, AAHR_1

### 3.2. Pharmacophore modelling/screening

The generated pharmacophore models were ranked automatically based on the survivality and BEDROC Score. The hypothesis AAHR_1 revealed the highest PhaseHypo Score of 0.837 comprising two hydrogen bond acceptors (A), one aromatic ring (R), and one Hydrophobic (H) features. The survival score of the hypothesis is 4.955, site score is 0.482, vector score is 0.847, volume score is 0.330, selectivity score is 1.283, and BEDROC score is 0.540.

The ∼3.5 million prepared ligand structures were screened using this AAHR_1 pharmacophore model and screened to 2,20,939 structures.

### 3.3. Pharmacophore Models Validation

The pharmacophore model developed was rigorously validated to assess its ability to accurately predict the activity of novel compounds identified through database screening or synthesized de novo. Validation is an essential step before utilizing a pharmacophore model for virtual screening (Kaserer *et al*., 2015).

The AAHR_1 model was applied to a test set database comprising 200 inactive molecules generated via the DUDE Decoys tool. It screened the decoy set to only 5 structures indicating an efficiency of 97.5%. Further, validation of the AAHR_1 hypothesis revealed that EF in the top 1% of the decoy dataset is 11.01%, demonstrating that pharmacophore model is 11.01-fold efficient in detecting true positives/actives from the entire dataset. ROC score, RIE, and AUAC values were calculated as 0.59, 5.29, and 0.68, respectively. Thus, AAHR_1 was statistically significant in picking the actives from the decoy dataset. Statistical significance of model was also validated by calculating BEDROC. Contrary to EF, BEDROC seeks to measure the early enrichment of the actives. BEDROC values were calculated at different tuning parameter values (α[=[8.0, α[=[20.0, and α[=[160.9) and found to be 0.520, 0.540, and 0.928, respectively.

### 3.4. Binding Affinity Prediction Analysis

The docking score range for the top 500 hits compounds was found between −15.588 to −9.701 kcal/mol, after XP docking. The top 2 compounds upon further screening based on MM-GBSA binding energy, i.e., CNP0375130 with MM-GBSA ΔG_bind_ of −76.41 kcal/mol and docking score of −13.567 kcal/mol, and CNP0256178 with MM-GBSA ΔG_bind_ of −75.02 kcal/mol and docking score of −14.027 kcal/mol, were selected for further studies.

The ligand CNP0375130 showed diverse interactions with multiple residues in the binding pocket of DNMT3A. It interacted with Arg891 with multiple H_2_O and an OH via H-bond and water bridges, Phe640 with H_2_O via H-bond, hydrophobic bond and salt bridge respectively. It also interacted with Arg836 through cation–π interaction of an aromatic ring via H-bond, hydrophobic bond and water bridges, and Arg887 via H-bonds and partially through water bridge with OH group. The ligand CNP0256178 showed interactions with multiple residues namely Arg790, Ser638, Gly706, Glu756 and Phe640 though different interactions like H-bond, hydrophobic bonds, water bridge, π-π stacking etc.

### 3.5. ADME/T Analysis

CNP0375130 and CNP0256178 exhibits moderate to good synthetic accessibility and comply with key drug-likeness rules, indicating favourable drug-like properties. While CNP0375130 meets criteria for drug-likeness, CNP0256178 adheres to Lipinski’s Rule of 5 and falls within the Golden Triangle and Pfizer Rule, suggesting good oral bioavailability potential. Despite moderate water solubility, both show promising human intestinal and oral absorption, with CNP0256178 demonstrating better permeability across membranes. They are metabolized by CYP3A4 with minimal CYP inhibition, reducing potential metabolic interactions. Both display a favorable safety profile with low acute toxicity, no mutagenicity, and no hERG inhibition. CNP0375130 exhibits a high maximum tolerated dose and low risk for carcinogenicity, skin sensitization, and respiratory toxicity. While CNP0256178 has a predicted moderate toxicity class, neither compound shows significant toxicity in various tests. Overall, the strengths of CNP0375130 and CNP0256178 suggests a promising candidate for further development. Addressing limitations like solubility and potential efflux challenges could lead to a more successful drug candidate. The in-depth details about the molecular, medicinal chemistry and ADMET properties of both the compounds are should in Table S1, S2 and S3 respectively.

### 3.6. Molecular Dynamics Simulation

#### 3.6.1. RMSD analysis

In Figure 5 (A), it is observed the backbone atoms of DNMT3A protein remained almost stable during the 100 ns simulation period for both the complexes A and B. For complex 5(A), the backbone of the protein slightly increased in the beginning till 28.62ns with an RMSD of 3.337 Å but later stabilized from till the end of the simulation with an average RMSD of 2.812 Å indicating that the protein reaches a relatively stable conformation. This stability is crucial as it suggests that the protein maintains its structural integrity over the simulation period. The initial increase in RMSD is common in MD simulations as the system equilibrates, but the eventual stabilization indicates a good quality simulation. The ligand’s RMSD increases rapidly within the first 10 ns, reaching around 10 Å, and fluctuates between 10-12 Å thereafter. This indicates that the ligand undergoes significant conformational changes or movements relative to the protein. The higher RMSD might suggest flexibility, which can be beneficial for the ligand to interact with different parts of the protein’s binding site.

**Figure 4.**
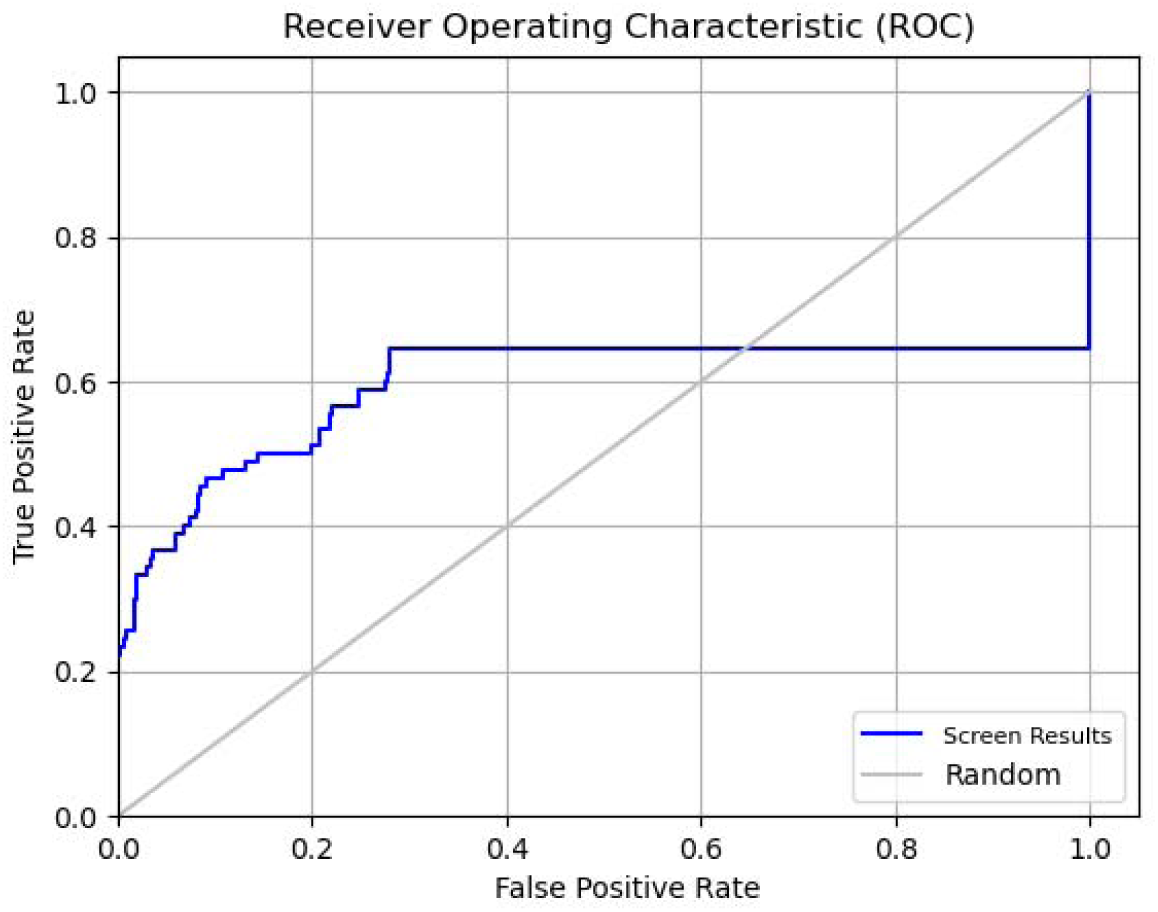
ROC curve represents the pharmacophore validation of the model AAHR_1

**Figure 5.**
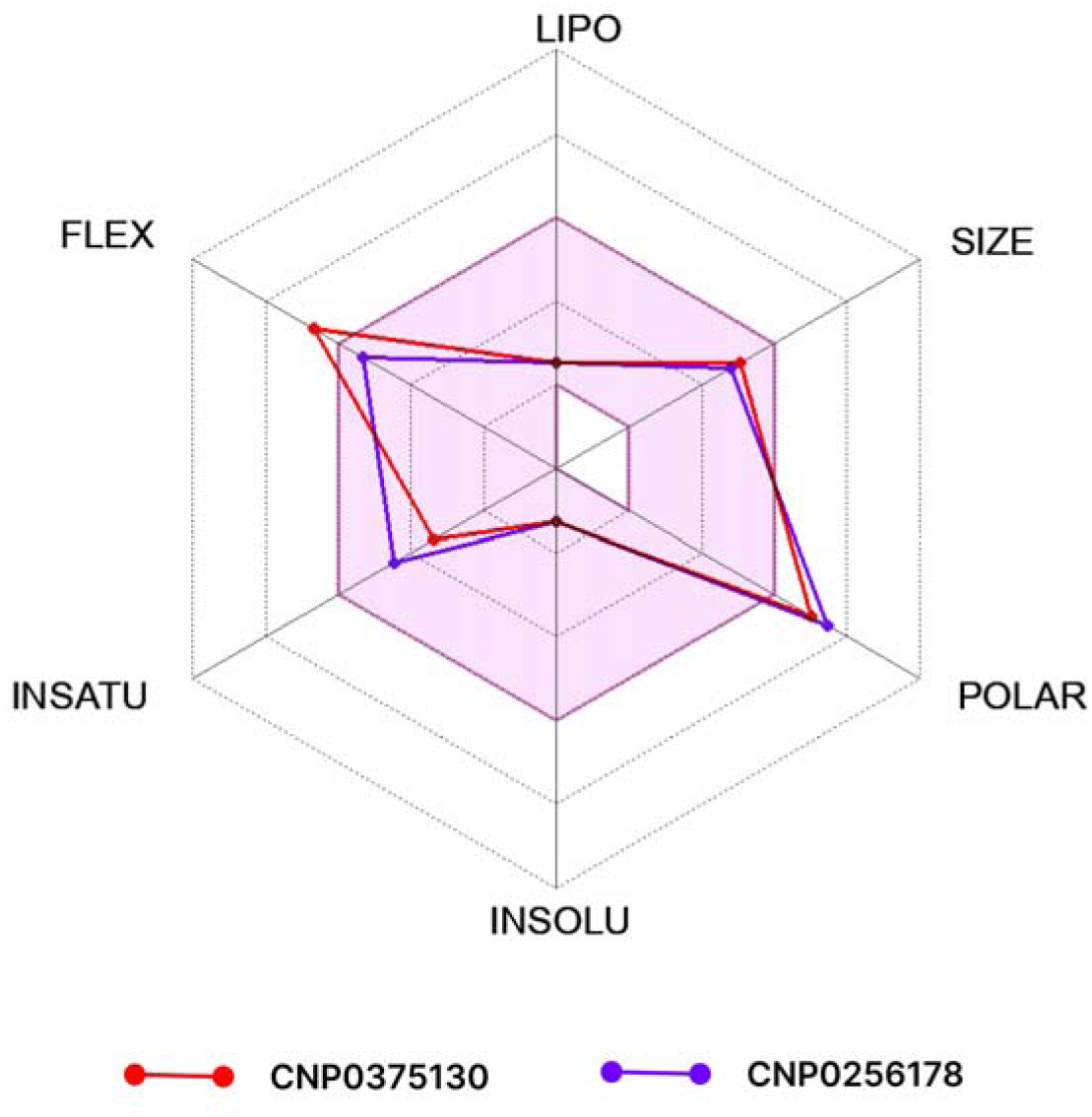
The bioavailability radar plot (obtained from SwissADME) depicting the excellent drug-likeliness of the two compounds (The reddish-brown area represents the optimal range for each property). The more detailed radar plot obtained from ADMETlab, are available in Figure S1 &S2

In Figure 5(B), The protein’s RMSD increases up to around 2.4 Å by 20 ns and then stabilizes between 2.4-2.8 Å, indicating a stable conformation is achieved relatively early in the simulation. This consistent stability is a positive indicator of the reliability of the simulation and the potential for the ligand to interact effectively without destabilizing the protein. The ligand’s RMSD increases rapidly in the first 10 ns, reaching around 3 Å, and then fluctuates between 3-5 Å. These values are lower compared to Graph A, indicating a more stable interaction with the protein. This relative stability suggests that the ligand maintains a consistent binding mode, which is crucial for effective drug design as it indicates strong and stable interactions with the protein’s active site.

#### 3.6.2. RMSF Analysis

The RMSF computations for the backbone atoms of 5YX2(chain A) was analysed (see Figure 6). The average RMSF values of the protein backbone was under 1.5Å for both the complexes. The average RMSF was found to be 1.37Å and 1.09Å for the complexes with ligands CNP0375130, and CNP0256178 respectively. Some slight fluctuations were seen at the residues interacting with the ligand atoms.

**Figure 6.**
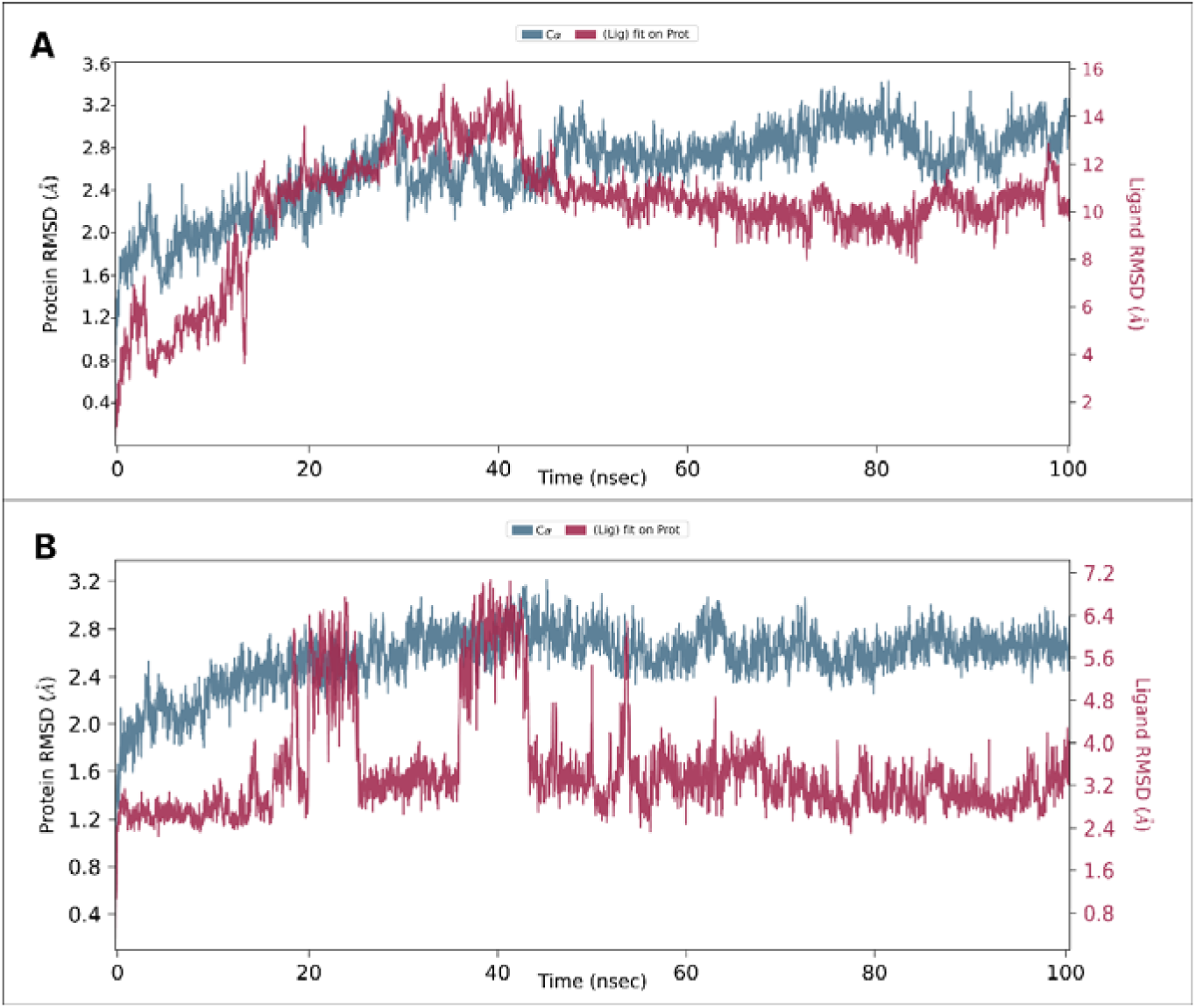
The line graph showing RMSD values of the complex structure extracted from ligand fit protein (ligand concerning protein) atoms. (A: 5YX2(A)-CNP0375130 and B: 5YX2(A)-CNP0256178)

RMSF values of the atoms of ligands CNP0375130 and CNP0256178 were computed to describe the dynamic behaviour of the ligands within the protein pocket (see Figure 7). The data indicate that some fluctuations in ligand CNP0375130 did not exceed 8.75Å for atom number 1, with an average of 5.2Å. For the ligand CNP0256178, the RMSF was maximum for atom number 26 with value of 4.49. The average RMSF values for the ligand was 1.81Å.

**Figure 7.**
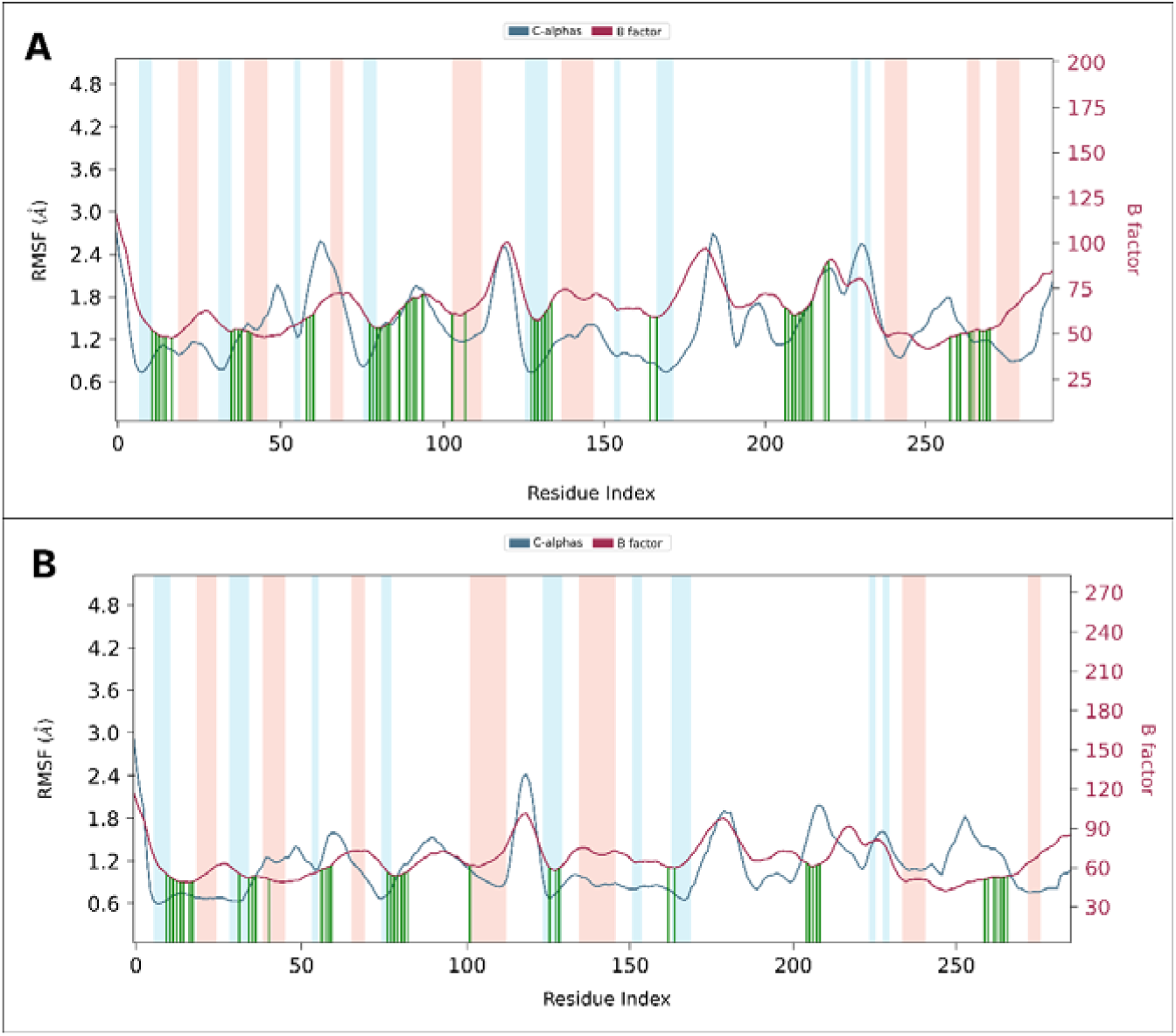
The line graph showing RMSF values of the complex structure extracted from protein residues backbone. The peaks of the blue line graph indicate the areas of the protein that fluctuate during the simulation. The maroon line graph indicates the B factor. The blue regions represent the alpha-helices and the orange region indicates the beta-helices. Protein residues that interact with the ligand are shown in green-coloured vertical lines. (A: 5YX2(A)-CNP0375130 and B: 5YX2(A)-CNP0256178.)

**Figure 8.**
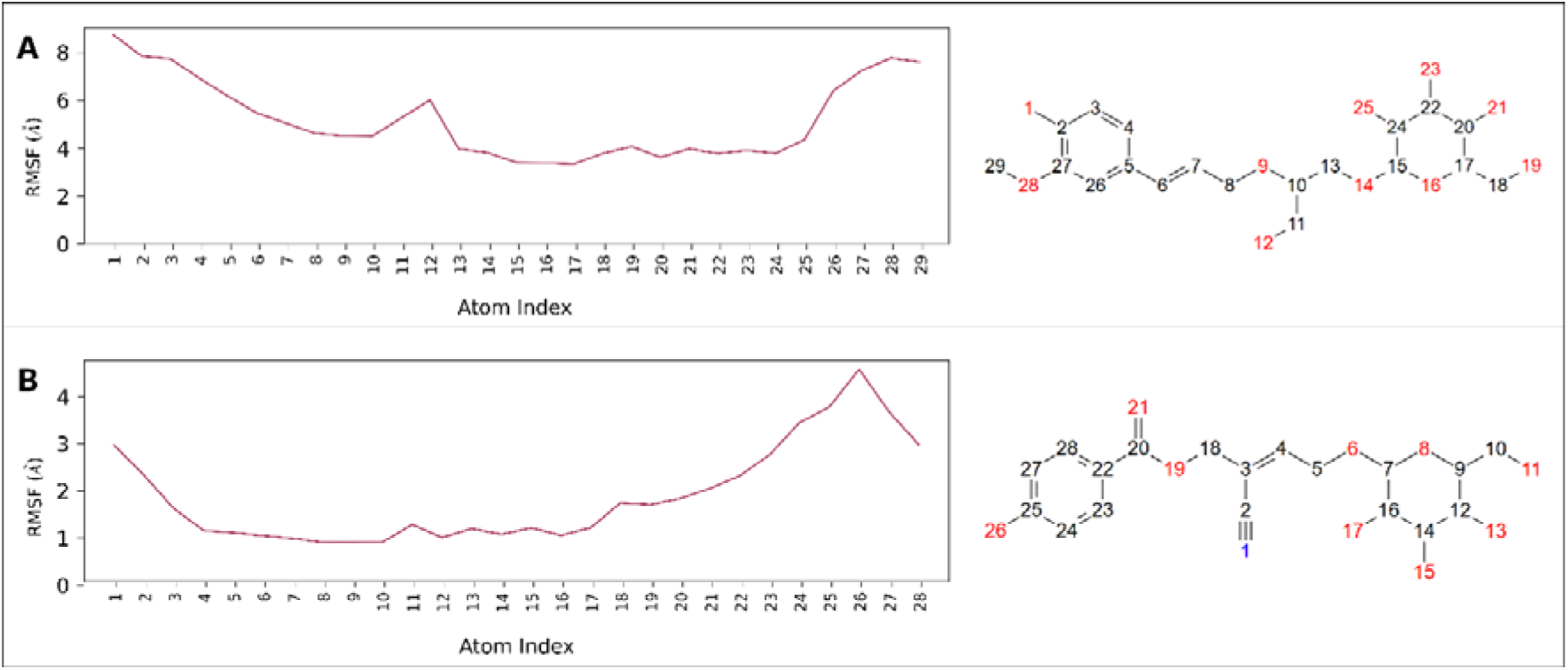
RMSF of Ligands’ atoms A: CNP0375130 and B: CNP0256178.

These slight fluctuations in ligand structures can be attributed to the flexible nature of the examined ligands. Nonetheless, the molecular structures of both ligands remained stable, wrt 5YX2(A) backbone atoms exhibiting stability over 100 ns in aqueous conditions.

#### 3.6.3. Protein-ligand contact dynamics

In the present study, it is identified four types of interactions within MD simulation: hydrogen bonds, hydrophobic interactions, water bridges, and ionic bonds. The timeline interactions of the residues of the protein with the ligands are shown in Figure 9, which shows a stable interaction.

**Figure 9.**
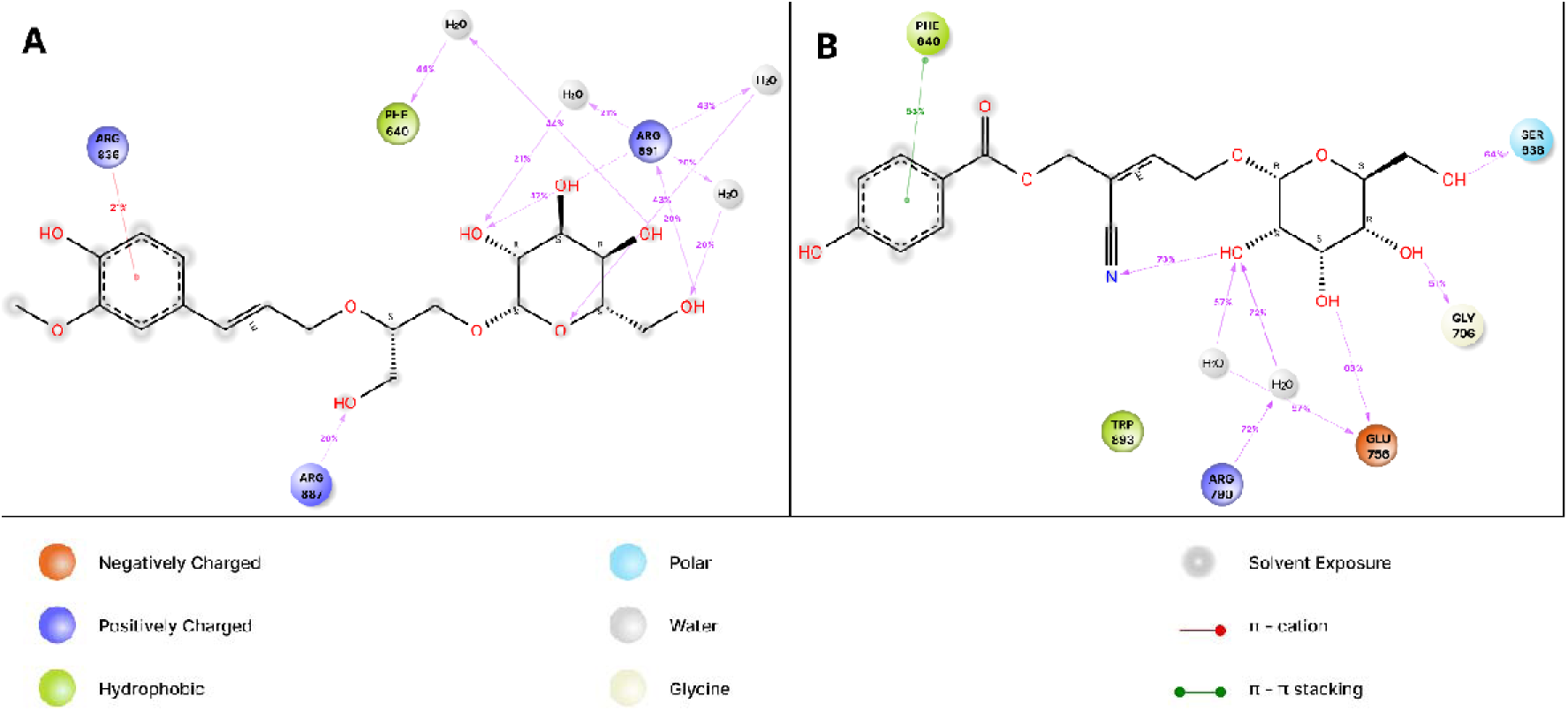
2D interaction diagram between protein-ligand complex A: 5YX2(A)-CNP0375130 and B: 5YX2(A)-CNP0256178.

The compound CNP0375130 showed significant hydrogen bond interactions with Agr891. Furthermore it also showed H-bond and water bridge interaction with Phe640, Arg792, Thr835 etc. CNP0256178 showed extensive H-bond interactions with residue Glu756. CNP0256178 showed additional interactions with amino acids such as Arg790, Phe640, Ser638 etc. majorly with H-bond, hydrophobic interactions and water bridges.

#### 3.6.4. Properties analysis of ligands CNP0375130 and CNP0256178

To validate the high stability of the examined complexes in an aqueous environment, we assessed the dynamic properties of ligands CNP0375130 and CNP0256178, which contribute to their stable interaction with the DNMT3A protein active sites. Figure S1(A,B) summarizes the six examined properties: Ligand RMSD, Radius of Gyration (rGyr), Molecular Surface Area (MolSA), Intramolecular Hydrogen Bonds (intraHB), Solvent Accessible Surface Area (SASA), and Polar Surface Area (PSA).

From Figure S1 (A,B), it is observed that the RMSD values of ligand CNP0375130, remained generally stable with an average of 2.07 Å. The RMSD values ranged between 0.45Å at 0.18 ns and 2.99Å at 29.1ns. On the other hand, the RMSD values of ligand CNP0256178 showed an average of 2.3Å throughout the 100 ns simulation, varying between 0.42Å at 0.12 ns and 2.83Å at 48.27 ns. The slight variations in RMSD values for all the two ligands (<∼3Å) during the 100 ns MD simulation indicate the high stability of ligands CNP0375130 and CNP0256178 in the protein pocket.

The average stability of rGyr values during the 100 ns MD simulation for CNP0375130 and CNP0256178 are 5.45Å and 5.05Å, respectively. For the properties MolSA, SASA, and PSA of ligands, the values were observed in the range of (361-399Å and 342-367Å), (55-429Å and 30-245Å), and (242-305Å and 250-326Å), with averages of 388.07 Å, 278.96 Å, 280.25 Å and 353.34 Å, 142.73 Å, 287.85 Å respectively. These findings further support the stability of ligands CNP0375130 and CNP0256178 in the 5YX2(A) protein environment.

### 3.7. Principal Component Analysis (PCA) and Dynamical Cross-Correlation Matrix (DCCM) Analysis

The PCA analysis (see Figure 10) of DNMT3A with ligand CNP0375130 (Graph A) and ligand CNP0256178 (Graph B) offers insightful comparisons into the conformational space and dynamics of these protein-ligand complexes. The PCA analysis of DNMT3A with ligands CNP0375130 (Graph A) and CNP0256178 (Graph B) reveals distinct impacts on protein-ligand dynamics. Graph A shows a compact, continuous distribution of points along PC1 and PC2, indicating that the DNMT3A-CNP0375130 complex occupies a confined conformational space with less variation in major collective motions. This suggests CNP0375130 binding stabilizes the protein, reducing flexibility and enhancing rigidity in key regions. The dense clustering implies a few dominant conformational states with frequent, closely related transitions, indicating stabilized dynamic behavior. Conversely, Graph B exhibits a broader, scattered distribution, indicating the DNMT3A-CNP0256178 complex samples a wider range of conformations, suggesting higher flexibility and a more extensive conformational space. Distinct clusters and discrete transitions suggest multiple conformational states, which could relate to different functional or interaction modes, indicating a more adaptable but potentially less stable structure compared to CNP0375130-bound form. The Internal PCA, Multi-Dimensional Scaling (MDS) and t-distributed Stochastic Neighbour Embedding (t-SNE) graphs are shown in figure S5.

**Figure 10.**
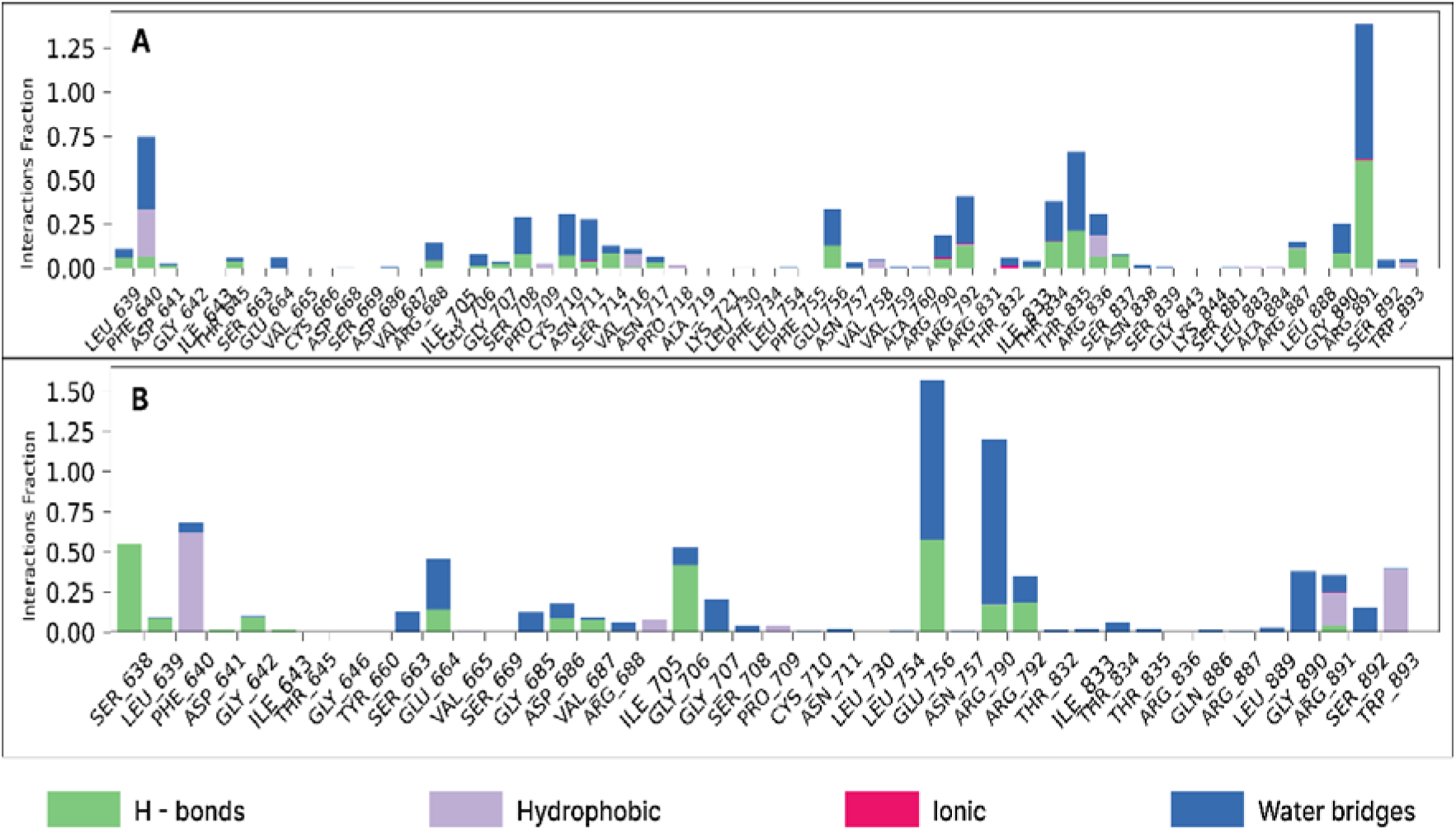
Interaction fraction histogram between protein-ligand complex A: 5YX2(A)-CNP0375130 and B: 5YX2(A)-CNP0256178.

The DCCM analysis (see Figure 11) of DNMT3A with ligands CNP0375130 (Graph A) and CNP0256178 (Graph B) reveals distinct residue correlated motions. Graph A shows strong positive correlations (red regions), indicating that CNP0375130 induces cooperative movements among residue clusters, likely stabilizing DNMT3A’s functional domains, critical for its functioning. Graph B displays dispersed positive and negative correlations (blue regions), suggesting that CNP0256178 causes heterogeneous, less coordinated movements, potentially leading to conformational changes and destabilization.

**Figure 11.**
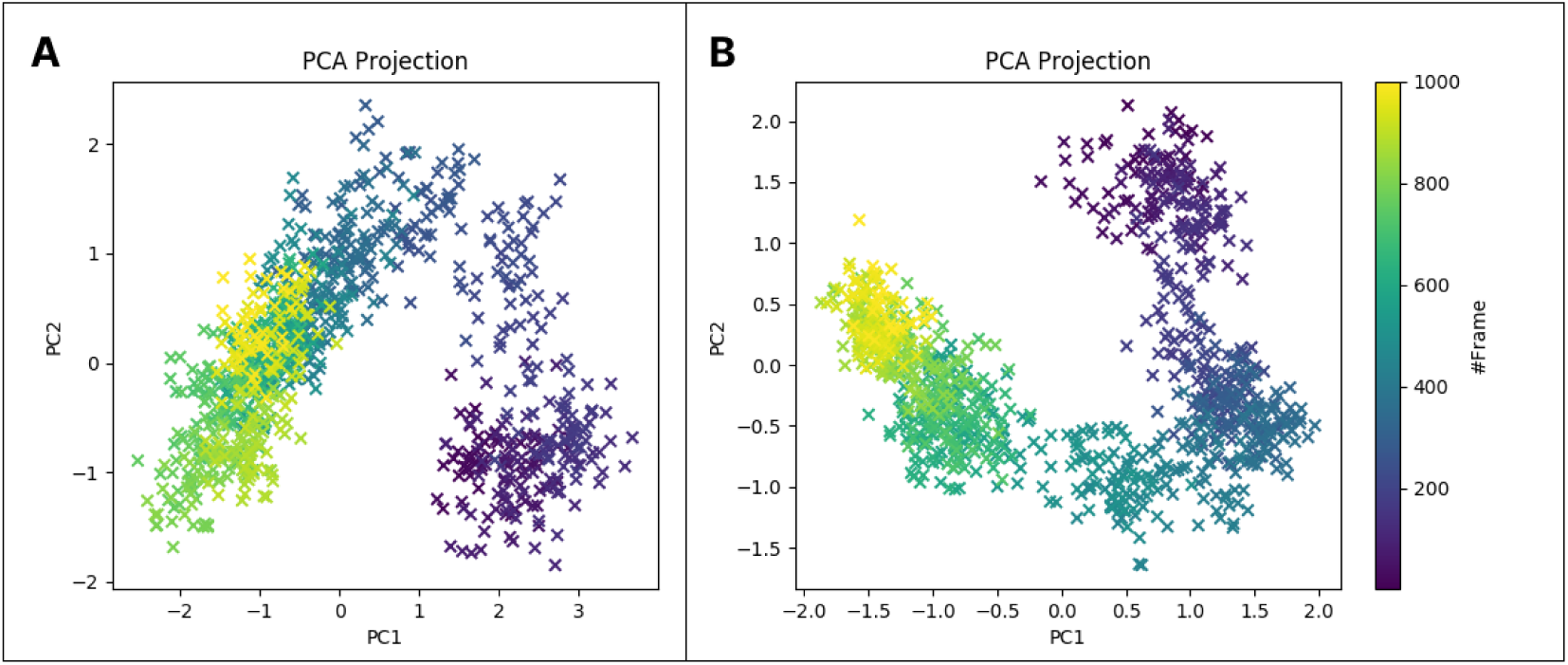
Standard Principal Component Analysis (PCA) graph of the trajectories of the two complexes A: 5YX2(A)-CNP0375130 and B: 5YX2(A)-CNP0256178.

**Figure 12.**
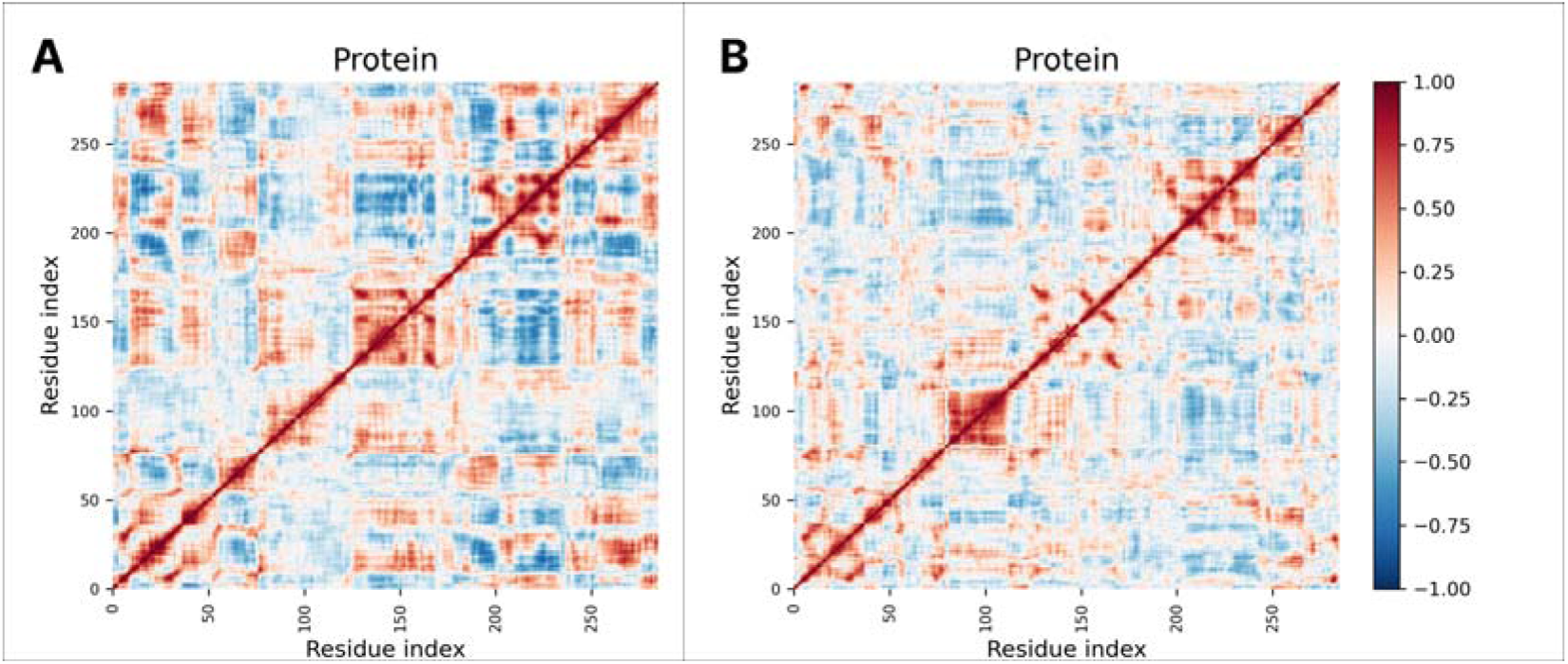
Dynamical Cross-Correlation Matrix (DCCM) Analysis graph of the trajectories of the two complexes A: 5YX2(A)-CNP0375130 and B: 5YX2(A)-CNP0256178. Red and blue colors indicate positive and negative correlations between residue pairs, respectively, while white represents minimal correlation. Strong diagonal dominance suggests local structural stability, with off-diagonal clusters indicating long-range interactions.

**Figure 13.**
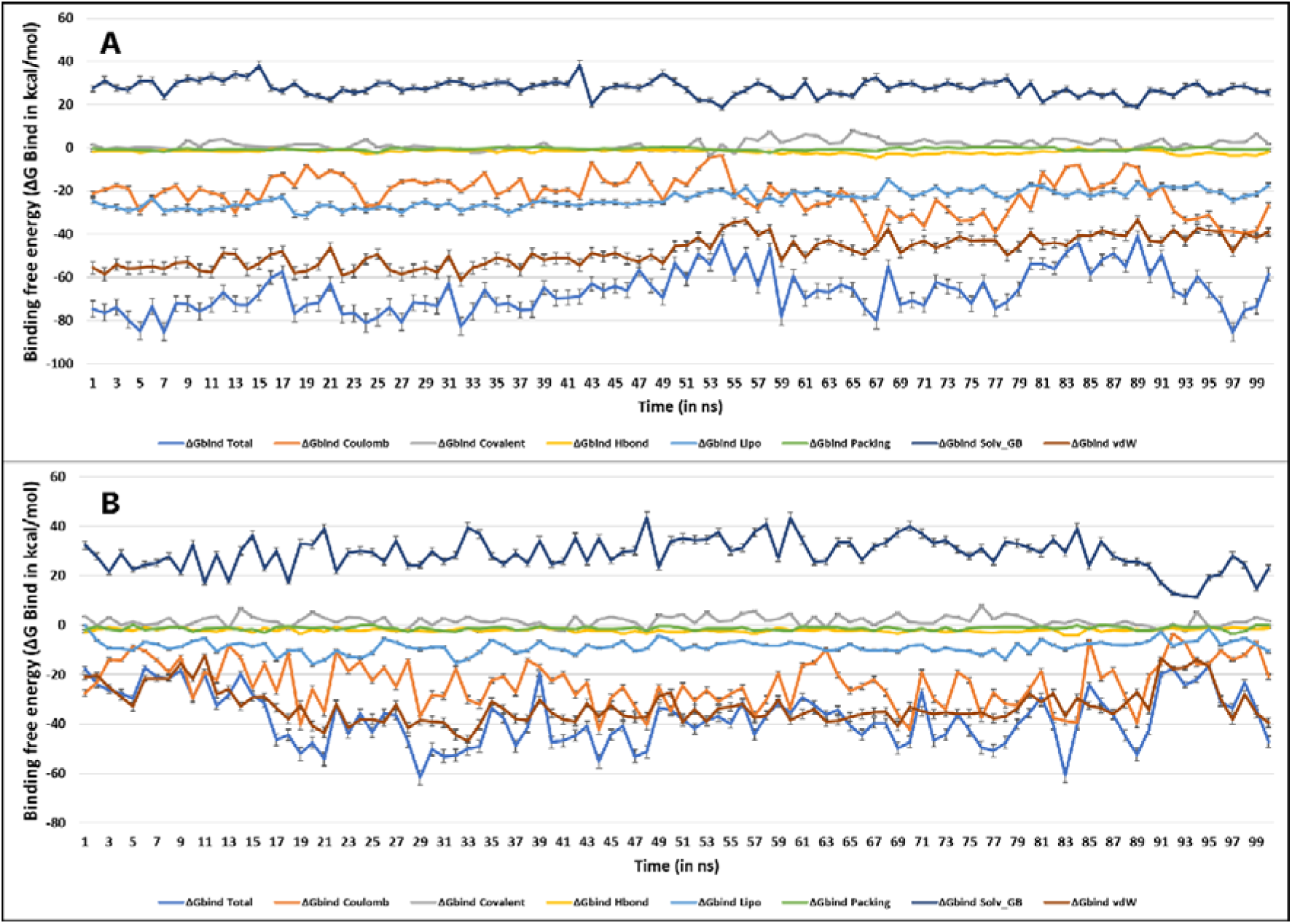
Post simulation MM-GBSA based Binding free energy (ΔG_Bind_ in kcal/mol) for (A) 5YX2(A)-CNP0375130 Complex; (B) 5YX2(A)-CNP0256178 Complex

Comparing the profiles, CNP0375130 fosters cohesive dynamic networks and structural stabilization in DNMT3A, enhancing its functional efficacy. Conversely, CNP0256178 introduces variability and flexibility, possibly affecting activity. These insights elucidate how each ligand uniquely modulates DNMT3A’s dynamics, explaining their distinct biological effects.

Integrating PCA with DCCM results, CNP0375130 promotes a stable, cohesive dynamic network in DNMT3A, with reduced flexibility (PCA) and enhanced correlated motions (DCCM). In contrast, CNP0256178 introduces greater flexibility and a broader conformational landscape, correlating with more heterogeneous, less coordinated residue motions. This suggests CNP0375130 is more effective in stabilizing DNMT3A’s functional conformation, while CNP0256178 allows for more dynamic states, potentially affecting the protein’s activity and interactions differently.

### 3.8. MM-GBSA Analysis

The parameters of free binding energies between the ligands were evaluated by the MM-GBSA simulations, which is the final step for checking the stability levels of the examined systems in the aqueous environment. The average values of calculated MM-GBSA energy parameters are presented in Table S4, containing binding free energies, Coulomb energy (Coulomb), Covalent bonding (Covalent), Hydrogen bonding (H-bond), lipophilic bonding (Lipo), π-π stacking interaction (Packing), solvent generalized bonding (SolvGB), and Van der Waals bonding energy (VDW) over the 100ns time frame.

From Table S4 and S5, the average ΔG_bind_ shown by 5YX2(A)-CNP0375130 Complex was −42.49 kcal/mol, maximum value of −76.01 kcal/mol, minimum of −23.14 kcal/mol, with a standard deviation (SD) of 8.41. The majority of it was contributed by ΔG_bind_ Coulomb (−19.94 kcal/mol), ΔG_bind_ Lipo (−12.24 kcal/mol) and, ΔGbind vdW (−35.35 kcal/mol). The average ΔG_bind_ shown by 5YX2(A)-CNP0256178 Complex was −55.25 kcal/mol, maximum value of −78.67 kcal/mol, minimum of −42.43 kcal/mol, with a standard deviation (SD) of 7.40. The majority of it was contributed by ΔG_bind_ Coulomb (−-29.25 kcal/mol), ΔG_bind_ Lipo (−17.58 kcal/mol) and, ΔG_bind_ vdW (−47.18 kcal/mol).

## 4. DISCUSSION

In this *in silico* study, the identified pharmacophore features align with previously reported binding characteristics of DNMT3A inhibitors (Yoo *et al*., 2012). CNP0375130 interacted with several key residues in the DNMT3A binding pocket, including Arg891, Phe640, Arg836, and Arg887 (Table 1). These interactions involved hydrogen bonds, water bridges, and cation-π interactions, suggesting a stable binding mode (Figure 3). CNP0256178 also interacted with multiple residues in the binding pocket, including Arg790, Ser638, Gly706, Glu756, and Phe640 (Table 1). The interactions involved hydrogen bonds, hydrophobic interactions, water bridges, and π-π stacking, suggesting a favorable binding mode (Figure 3). CNP0375130, classified as a lipid and lipid-like molecule under glycerolipids, specifically glycosylglycerols, was obtained from the Supernatural3 database according to the CONONUT database. CNP0256178, also a lipid and lipid-like molecule, belongs to fatty acyls and fatty acyl glycosides of mono- and disaccharides, and was sourced from UNPD (Universal Natural Products Database) and Super Natural II. ADMET analysis predicted favorable drug-like properties, bioavailability, and safety profiles for both CNP0375130 and CNP0256178. MD simulations revealed stable interactions between the ligands and DNMT3A protein. CNP0375130 formed more stable interactions with the protein compared to CNP0256178, as evidenced by lower ligand RMSD values (Figure 5). PCA and DCCM analyses again suggested that CNP0375130 induced a more rigid and cohesive dynamic network in DNMT3A, potentially enhancing protein stability (Figures 10 & 11). The identified binding residues suggest that both CNP0375130 and CNP0256178 interact with a combination of hydrophobic and electrostatic interactions within the DNMT3A binding pocket. The presence of multiple interacting residues, particularly arginines (positively charged) for CNP0375130, suggests potential for strong binding affinity. Arginines within the DNMT3A active site are crucial for substrate (DNA) binding and catalysis (Cull *et al*., 2018). Their positively charged side chains can form strong salt bridges with negatively charged phosphates on the DNA backbone, facilitating proper substrate positioning for enzymatic activity (Yusufaly *et al*., 2014). Studies have shown hydrophobic residues like phenylalanine contribute to the formation of hydrophobic pockets within the DNMT3A active site (Jeltsch & Jurkowska, 2016). These pockets are essential for accommodating specific functional groups on the substrate or inhibitor, promoting favorable interactions and binding specificity (Patil *et al*., 2010). Depending on their ionization state (pH-dependent), serine and glutamate residues can participate in hydrogen bonding with the inhibitor or substrate. Hydrogen bonds are crucial for fine-tuning binding affinity and stabilizing protein-ligand complexes. The presence of multiple arginines (Arg891, Arg887, potentially Arg836) interacting with CNP0375130 aligns with the importance of these residues for substrate binding in DNMT3A, suggesting CNP0375130 might competitively bind to the same site as the DNA substrate, potentially mimicking its interactions and effectively inhibiting enzyme activity. The interaction with Phe640 in both ligands suggests this residue creates a hydrophobic pocket within the binding site that is crucial for ligand accommodation, targeting this pocket with well-designed functional groups on the inhibitors could enhance binding affinity and specificity. The interaction with Glu756 is significant as Glu756 is known to be a critical residue for inhibitor binding in DNMT3A inhibitors. Glu756, located in the active site of DNMT3A, plays a crucial role in the enzyme’s catalytic activity and substrate recognition. Many DNMT3A inhibitors are designed to interact with the catalytic site residues, including Glu756, to effectively inhibit the enzyme’s activity (W. Yang *et al*., 2023). The interaction with Glu756 can significantly impact the binding affinity and inhibitory potency of the compounds. In various studies, mutations or modifications at or near Glu756 have been shown to affect the binding and efficacy of inhibitors, highlighting its importance in the structure-function relationship of DNMT3A (Kudo *et al*., 2024; Xie *et al*., 2019). Therefore, targeting Glu756 is a common strategy in the design of DNMT3A inhibitors for therapeutic purposes. Based on the promising *in silico* results, further research is recommended, including *in vitro* and *in vivo* studies to evaluate the anti-leukemic activity of CNP0375130 and CNP0256178 in AML cell lines and animal models, mechanism of action studies to experimentally determine the specific mechanism by which these compounds inhibit DNMT3A activity, selectivity studies to assess the selectivity of the compounds towards DNMT3A compared to other proteins to minimize potential off-target effects, and structure-activity relationship (SAR) studies to investigate how modifications to the identified ligand structures can improve their potency and selectivity.

**Table 1.** Top 2 compounds with COCONUT ID, molecular formula, top interacting residues, and XP docking score (kcal/mol) and MM-GBSA ΔG_bind_ energy (kcal/mol)

| Compound | Formula | Major Interacting Residues | Docking Score | MM-GBSA $\Delta G_{\text{bind}}$ |
| --- | --- | --- | --- | --- |
| **CNP0375130** | C<sub>19</sub>H<sub>28</sub>O<sub>10</sub> | F640, R891, R887, R836 | -13.567 | -76.41 |
| **CNP0256178** | C<sub>18</sub>H<sub>21</sub>NO<sub>9</sub> | R790, G706, E756, S638, F640 | -14.027 | -75.02 |

## 5. CONCLUSION

In this research, we utilized advanced computational methodologies to identify novel inhibitors of DNMT3A, a pivotal epigenetic regulator in acute myeloid leukemia (AML). Through a synergistic application of pharmacophore modeling, virtual screening, and molecular dynamics simulations, we identified CNP0375130 and CNP0256178 as promising lead compounds. These candidates, derived from natural compounds, exhibited robust binding affinities and favorable physicochemical properties, positioning them as viable drug candidates. This study highlights the efficacy of computational drug discovery approaches in accelerating the identification of novel therapeutics for complex diseases such as leukemia. The lead compounds identified here offer a solid foundation for future optimization and *in vitro* and *in vivo* studies, marking a critical step towards the development of effective treatments for AML. Additionally, the use of natural compounds underscores the potential of harnessing bioactive molecules from nature in the quest for innovative drug therapies.

## Supporting information

Supplimentary_DNMT3A

