## Supplementary material for "*In silico* Exploration Natural Compounds for the Discovery of Novel DNMT3A Inhibitors as Potential Therapeutic Agents for Acute Myeloid Leukemia": Supplimentary_DNMT3A

Table 1 Physiochemical properties of the two screened compounds

| Properties | CNP0375130 | CNP0256178 |
| --- | --- | --- |
| Molecular Weight | 416.17 | 395.120 |
| Dipole Moment | 3.132 | 8.835 |
| Density | 1.047 | 1.055 |
| H-bond donors | 6 | 5 |
| H-bond acceptors | 10 | 10 |
| Number of rotatable bonds | 10 | 8 |
| Number of rings | 2 | 2 |
| Number of atoms in the biggest ring | 6 | 6 |
| Number of heteroatoms | 10 | 10 |
| Formal Charge | 0 | 0 |
| Number of rigid bonds | 13 | 15 |
| Sterio Centres | 6 | 5 |
| SASA | 714.245 | 672.398 |
| FOSA | 327.154 | 199.252 |
| FISA | 271.482 | 302.037 |
| PISA | 115.609 | 171.109 |
| WPSA | 0 | 0 |
| Total solvent accessible volume (in Å^3^) | 1269.31 | 1190.78 |
| Polarizability (in Å^3^) | 35.474 | 34.788 |
| Hexadecane/Gas partition coefficient | 14.158 | 13.761 |
| Octanol/Gas partition coefficient | 28.582 | 27.709 |
| Water/Gas partition coefficient | 24.202 | 23.244 |
| Octanol/Water partition coefficient | -1.023 | -1.231 |
| Number of non-conjugated amine groups | 0 | 0 |
| Number of amidine and guanidine groups | 0 | 0 |
| Number of carboxylic acid groups | 0 | 0 |
| Number of non-conjugated amide groups | 0 | 0 |
| Number of reactive functional groups | 1 | 2 |
| PM3 calculated ionization potential (negative of HOMO energy) (in eV) | 8.86 | 9.754 |
| PM3 calculated electron affinity (negative of LUMO energy) (in eV) | 0.365 | 0.616 |

Table 2 Medicinal Chemistry properties of the two screened compounds

| Properties | CNP0375130 | CNP0256178 |
| --- | --- | --- |
| Synthetic Accessibility Score | 5.13 | 4.78 |
| MCE-18 Value | 50.00 | 50.769 |
| PAINS | 0 alert | 0 alert |
| Brenk | 0 alert | 1 alert |
| ALARM NMR Rule | 2 alert | 2 alert |
| BMS Rule | 0 alert | 0 alert |
| Chelator Rule | 1 alert | 0 alert |
| Lipinski’s Rule of 5 violations | 0 violations | 0 violations |
| Golden Triangle | Accepted | Accepted |
| Pfizer Rule | Accepted | Accepted |

Table 3 ADMET properties of the two screened compounds

| **Properties** | | **CNP0375130** | **CNP0256178** |
| --- | --- | --- | --- |
| **A** | Water solubility (log mol/L) | -4.434 | -4.646 |
|  | Caco-2 permeability (Papp in 10^-6^ cm/s) | 0.336 | 0.329 |
|  | Human intestinal absorption (in %) | 67.464 | 69.605 |
|  | Skin Permeability (log K_p_) | -2.909 | -2.954 |
|  | Human oral absorption (in %) | 73.436 | 79.991 |
|  | P-glycoprotein substrate | Yes | Yes |
|  | P-glycoprotein I inhibitor | No | No |
|  | P-glycoprotein II inhibitor | No | No |
|  | MDCK Permeability (in nm/sec) | 0.00049 | 0.00011 |
| **D** | Volume of distribution at steady state (in log L/kg) | -0.331 | -0.391 |
|  | Plasma protein binding (in %) | 39.426 | 54.459 |
|  | Fraction of unbound plasms | 0.513 | 0.578 |
|  | BBB Permeability (in logBB) | -1.057 | -1.009 |
|  | CNS permeability (in logPS) | -3.271 | -3.23 |
| **M** | CYP2D6 substrate | No | No |
|  | CYP3A4 substrate | Yes | Yes |
|  | CYPIA2 inhibitor | No | No |
|  | CYP2C19 inhibitor | No | No |
|  | CYP2C9 inhibitor | No | No |
|  | CYP2D6 inhibitor | No | No |
|  | CYP3A4 inhibitor | No | No |
| **E** | Total Clearance (in log ml/min/kg) | 1.526 | 1.561 |
|  | Renal OCT2 substrate | No | No |
| **T** | Predicted GHS Toxicity Class | 5 | 5 |
|  | AMES toxicity | No | No |
|  | Max. tolerated dose (human) | -0.369 | -0307 |
|  | hERG I inhibitor | No | No |
|  | hERG II inhibitor | No | No |
|  | Oral Rat Acute Toxicity (LD50) (mol/kg) | 2.89 | 3.08 |
|  | Oral Rat Chronic Toxicity (LOAEL) (log mg/kg/day) | 1.078 | 1.023 |
|  | Skin Sensitisation | No | No |
|  | *T Pyriformis* toxicity (µg/L) | 0.571 | 0.574 |
|  | Minnow toxicity (log mM) | 2.638 | 2.795 |
|  | DILI | No | No |
|  | FDA Maximum Recommended Daily Dose | 0.005 | 0.036 |
|  | Carcinogenicity | No | No |
|  | Eye corrosion | No | No |
|  | Eye irritation | No | No |
|  | Respiratory toxicity | No | No |

Table 4 Post simulation MM-GBSA based Binding free energy (ΔG_Bind_ in kcal/mol) for the 5YX2(A)- CNP0375130 Complex

| Time (in ns) | ΔG_bind_ Total | ΔG_bind_ Coulomb | ΔG_bind_ Covalent | ΔG_bind_ Hbond | ΔG_bind_ Lipo | ΔG_bind_ Packing | ΔG_bind_ Solv_GB | ΔG_bind_ vdW |
| --- | --- | --- | --- | --- | --- | --- | --- | --- |
| 1 | -76.01 | -35.95 | 6.42 | -4.29 | -26.16 | -0.79 | 39.31 | -54.56 |
| 2 | -63.46 | -25.30 | 3.25 | -2.32 | -21.02 | -2.04 | 32.25 | -48.29 |
| 3 | -57.18 | -21.40 | -1.84 | -2.46 | -16.19 | -1.59 | 27.06 | -40.75 |
| 4 | -48.13 | -22.73 | 0.67 | -3.45 | -16.31 | -0.08 | 35.38 | -41.61 |
| 5 | -55.46 | -25.43 | 3.49 | -1.48 | -20.05 | -1.87 | 39.42 | -49.54 |
| 6 | -54.69 | -35.84 | 4.50 | -3.19 | -17.49 | -2.31 | 37.31 | -37.67 |
| 7 | -47.70 | -27.54 | 1.22 | -2.22 | -14.52 | -2.30 | 36.38 | -38.73 |
| 8 | -48.77 | -19.70 | 4.93 | -1.76 | -14.16 | -1.37 | 24.10 | -40.80 |
| 9 | -61.65 | -24.33 | 5.29 | -2.69 | -18.48 | -1.35 | 31.59 | -51.68 |
| 10 | -65.20 | -21.52 | 3.32 | -1.79 | -22.23 | -0.77 | 33.94 | -56.15 |
| 11 | -59.09 | -19.42 | 1.21 | -2.64 | -15.82 | -0.52 | 24.64 | -46.53 |
| 12 | -51.79 | -25.52 | 0.37 | -2.44 | -12.44 | -1.77 | 32.42 | -42.40 |
| 13 | -38.81 | -15.12 | 2.34 | -1.13 | -12.22 | -0.02 | 23.76 | -36.43 |
| 14 | -36.67 | -18.67 | 1.28 | -2.57 | -12.28 | -0.03 | 23.03 | -27.43 |
| 15 | -34.12 | -24.75 | 1.02 | -3.27 | -9.78 | 0.00 | 28.08 | -25.42 |
| 16 | -39.43 | -25.27 | 2.21 | -3.88 | -10.49 | -0.55 | 28.26 | -29.71 |
| 17 | -27.14 | -13.91 | 2.13 | -1.97 | -10.31 | -0.40 | 26.95 | -29.64 |
| 18 | -35.62 | -30.29 | 4.05 | -2.19 | -11.47 | -1.17 | 41.63 | -36.18 |
| 19 | -23.14 | -13.29 | 4.07 | -1.15 | -7.46 | -1.31 | 29.42 | -33.42 |
| 20 | -36.74 | -33.15 | 4.62 | -2.25 | -10.54 | -0.99 | 36.47 | -30.90 |
| 21 | -40.27 | -17.16 | -0.22 | -1.24 | -9.24 | -0.11 | 20.51 | -32.82 |
| 22 | -34.60 | -29.16 | 1.11 | -3.30 | -7.35 | -0.09 | 33.18 | -28.99 |
| 23 | -30.44 | -12.13 | 2.22 | -1.34 | -7.80 | -0.82 | 23.71 | -34.27 |
| 24 | -46.26 | -36.66 | 3.37 | -2.62 | -10.25 | -0.29 | 27.24 | -27.04 |
| 25 | -47.98 | -38.92 | 3.92 | -2.57 | -9.14 | -0.56 | 30.42 | -31.13 |
| 26 | -35.83 | -33.32 | 4.64 | -3.84 | -7.66 | -0.12 | 28.76 | -24.29 |
| 27 | -42.03 | -19.61 | 1.03 | -3.27 | -10.66 | -0.02 | 20.47 | -29.98 |
| 28 | -36.93 | -18.59 | -1.20 | -3.19 | -9.58 | -0.17 | 24.78 | -28.98 |
| 29 | -32.41 | -12.28 | 1.21 | -0.97 | -10.23 | -0.51 | 22.36 | -31.98 |
| 30 | -41.90 | -28.08 | 9.35 | -3.74 | -11.24 | -0.13 | 24.65 | -32.71 |
| 31 | -47.21 | -20.22 | 4.86 | -2.24 | -10.03 | -1.09 | 14.41 | -32.90 |
| 32 | -37.82 | -14.58 | 0.32 | -1.81 | -6.40 | -0.35 | 18.58 | -33.56 |
| 33 | -37.81 | -5.16 | 3.94 | -0.67 | -11.00 | -1.17 | 15.22 | -38.98 |
| 34 | -43.61 | -32.04 | 7.64 | -2.22 | -10.23 | -2.24 | 28.47 | -32.98 |
| 35 | -53.32 | -29.60 | 2.95 | -2.14 | -11.71 | -1.40 | 21.66 | -33.07 |
| 36 | -50.64 | -30.08 | 4.97 | -3.73 | -10.80 | -1.79 | 26.20 | -35.40 |
| 37 | -36.86 | -23.32 | 4.21 | -3.75 | -10.76 | -0.81 | 28.65 | -31.08 |
| 38 | -42.07 | -14.49 | 2.02 | -2.38 | -10.57 | -1.89 | 21.28 | -36.03 |
| 39 | -45.66 | -17.17 | 3.42 | -3.80 | -12.09 | -1.40 | 25.37 | -39.99 |
| 40 | -47.02 | -27.33 | 5.19 | -2.62 | -11.30 | -1.58 | 27.77 | -37.15 |
| 41 | -50.94 | -34.46 | 4.05 | -3.69 | -9.71 | -1.44 | 26.49 | -32.18 |
| 42 | -39.46 | -12.77 | 2.29 | -2.48 | -10.68 | -1.54 | 24.07 | -38.35 |
| 43 | -48.01 | -20.99 | 4.87 | -2.19 | -10.97 | -1.90 | 19.22 | -36.07 |
| 44 | -37.19 | -11.30 | 1.24 | -1.42 | -10.77 | -1.82 | 25.45 | -38.57 |
| 45 | -43.49 | -27.48 | 5.33 | -1.04 | -12.55 | -1.96 | 32.79 | -38.57 |
| 46 | -48.71 | -25.64 | 2.79 | -2.82 | -9.79 | -2.40 | 21.87 | -32.73 |
| 47 | -39.22 | -22.66 | 2.28 | -2.17 | -7.88 | -0.72 | 24.72 | -32.79 |
| 48 | -38.68 | -32.43 | 2.04 | -2.57 | -7.51 | -0.84 | 36.99 | -34.36 |
| 49 | -39.99 | -20.05 | 6.58 | -1.73 | -12.42 | -1.30 | 31.30 | -42.37 |
| 50 | -32.43 | -7.85 | -1.06 | -1.07 | -7.51 | -1.79 | 21.28 | -34.43 |
| 51 | -43.43 | -9.56 | 2.21 | -1.35 | -15.77 | 0.00 | 24.39 | -43.34 |
| 52 | -35.68 | -12.35 | 1.76 | -0.95 | -10.97 | -1.43 | 24.49 | -36.23 |
| 53 | -46.78 | -18.72 | 2.89 | -1.12 | -13.10 | -1.28 | 28.09 | -43.53 |
| 54 | -36.60 | -16.81 | 2.74 | -1.08 | -10.85 | -0.96 | 23.44 | -33.07 |
| 55 | -42.24 | -22.95 | 3.28 | -2.49 | -10.68 | -3.21 | 26.57 | -32.77 |
| 56 | -42.57 | -16.78 | 5.56 | -1.69 | -12.76 | -3.63 | 22.20 | -35.48 |
| 57 | -43.88 | -15.25 | 3.43 | -2.40 | -12.46 | -0.84 | 20.33 | -36.70 |
| 58 | -41.07 | -15.37 | 3.42 | -1.55 | -12.22 | -2.57 | 24.41 | -37.19 |
| 59 | -40.60 | -19.89 | 3.32 | -2.85 | -11.75 | -1.23 | 23.19 | -31.38 |
| 60 | -42.63 | -12.41 | 0.76 | -1.62 | -11.18 | -1.92 | 21.51 | -37.77 |
| 61 | -45.14 | -19.50 | 2.11 | -1.76 | -12.51 | -3.62 | 29.50 | -39.35 |
| 62 | -44.57 | -24.63 | 5.86 | -2.93 | -11.91 | -2.07 | 21.49 | -30.38 |
| 63 | -44.03 | -17.88 | 2.42 | -1.62 | -14.69 | -2.59 | 25.94 | -35.60 |
| 64 | -33.85 | -10.10 | 3.91 | -1.00 | -13.80 | -0.37 | 20.27 | -32.76 |
| 65 | -36.59 | -22.17 | 3.75 | -1.90 | -7.41 | -0.25 | 25.75 | -34.36 |
| 66 | -36.39 | -15.28 | 3.58 | -1.85 | -10.18 | -0.76 | 16.38 | -28.28 |
| 67 | -37.94 | -15.86 | 3.11 | -1.46 | -10.06 | -3.13 | 24.19 | -34.72 |
| 68 | -35.77 | -10.98 | 4.73 | -1.53 | -10.90 | -3.13 | 19.21 | -33.15 |
| 69 | -33.20 | -10.37 | 2.44 | -0.89 | -9.71 | -2.96 | 15.68 | -27.39 |
| 70 | -43.95 | -12.56 | -3.16 | -1.47 | -8.93 | -3.68 | 20.30 | -34.44 |
| 71 | -43.20 | -21.71 | 3.71 | -1.76 | -12.42 | -3.49 | 26.45 | -33.98 |
| 72 | -36.85 | -14.06 | 4.79 | -0.70 | -11.79 | -2.95 | 23.56 | -35.71 |
| 73 | -40.90 | -16.91 | 3.14 | -0.86 | -12.85 | -1.51 | 24.51 | -36.41 |
| 74 | -39.52 | -3.73 | 0.65 | -0.60 | -13.66 | -0.72 | 20.86 | -42.31 |
| 75 | -50.29 | -21.34 | 5.88 | -3.16 | -14.67 | -1.69 | 21.02 | -36.33 |
| 76 | -38.78 | -11.33 | 5.01 | -1.65 | -14.86 | -2.34 | 22.52 | -36.14 |
| 77 | -28.71 | -7.39 | 3.00 | -0.71 | -9.60 | -2.74 | 19.23 | -30.50 |
| 78 | -34.54 | -11.54 | 1.46 | -0.74 | -10.66 | -3.16 | 22.36 | -32.25 |
| 79 | -32.58 | -13.44 | 4.36 | -0.95 | -11.39 | -2.66 | 26.08 | -34.57 |
| 80 | -37.75 | -26.46 | 4.40 | -1.95 | -10.32 | -1.94 | 28.95 | -30.42 |
| 81 | -34.49 | -18.83 | 7.74 | -0.80 | -11.73 | -2.54 | 26.39 | -34.72 |
| 82 | -36.43 | -6.92 | 2.25 | -0.46 | -12.22 | -2.40 | 20.40 | -37.09 |
| 83 | -37.20 | -10.85 | 1.25 | -1.57 | -11.76 | -2.85 | 25.52 | -36.94 |
| 84 | -36.94 | -17.90 | 3.50 | -1.00 | -11.55 | -3.13 | 25.56 | -32.42 |
| 85 | -31.45 | -7.04 | 3.31 | -1.05 | -10.47 | -2.54 | 16.10 | -29.76 |
| 86 | -43.29 | -18.57 | 2.67 | -1.89 | -12.90 | -0.36 | 23.26 | -35.49 |
| 87 | -46.21 | -34.80 | 3.75 | -4.19 | -13.25 | -0.06 | 29.96 | -27.63 |
| 88 | -45.18 | -20.43 | 1.88 | -2.25 | -13.29 | -0.20 | 19.85 | -30.74 |
| 89 | -44.61 | -19.20 | 0.94 | -1.83 | -14.05 | 0.00 | 23.16 | -33.62 |
| 90 | -34.64 | -9.49 | 2.29 | -0.74 | -12.50 | -1.37 | 21.06 | -33.87 |
| 91 | -44.27 | -21.09 | 3.28 | -2.47 | -15.01 | -1.25 | 28.66 | -36.39 |
| 92 | -45.13 | -22.80 | 2.91 | -2.53 | -14.67 | -0.39 | 23.87 | -31.51 |
| 93 | -51.20 | -23.57 | 4.16 | -1.97 | -16.62 | -0.07 | 22.21 | -35.35 |
| 94 | -53.64 | -31.90 | 6.84 | -2.42 | -15.23 | -1.44 | 26.57 | -36.06 |
| 95 | -41.80 | -22.52 | 7.77 | -2.46 | -13.48 | -0.65 | 22.37 | -32.81 |
| 96 | -41.55 | -16.46 | 6.53 | -1.97 | -14.88 | 0.00 | 19.42 | -34.19 |
| 97 | -47.09 | -18.46 | 2.78 | -1.74 | -15.85 | 0.00 | 22.11 | -35.94 |
| 98 | -34.23 | -11.13 | 8.51 | -1.80 | -14.51 | -0.47 | 17.33 | -32.16 |
| 99 | -49.57 | -17.09 | 4.54 | -3.07 | -13.80 | 0.00 | 13.76 | -33.91 |
| 100 | -50.11 | -22.81 | 2.71 | -1.50 | -14.65 | -0.90 | 23.72 | -36.68 |
| AVG | **-42.49** | **-19.94** | **3.24** | **-2.06** | **-12.24** | **-1.37** | **25.23** | **-35.35** |
| MAX | **-76.01** | **-38.92** | **-3.16** | **-4.29** | **-26.16** | **-3.68** | **13.76** | **-56.15** |
| MIN | **-23.14** | **-3.73** | **9.35** | **-0.46** | **-6.40** | **0.00** | **41.63** | **-24.29** |
| SD | **8.41** | **7.84** | **2.15** | **0.92** | **3.22** | **1.04** | **5.63** | **5.62** |

Table 5 Post simulation MM-GBSA based Binding free energy (ΔGBind in kcal/mol) for the 5YX2(A)- CNP0256178 Complex

| Time (in ns) | ΔG_bind_ Total | ΔG_bind_ Coulomb | ΔG_bind_ Covalent | ΔG_bind_ Hbond | ΔG_bind_ Lipo | ΔG_bind_ Packing | ΔG_bind_ Solv_GB | ΔG_bind_ vdW |
| --- | --- | --- | --- | --- | --- | --- | --- | --- |
| 1 | -66.25 | -23.24 | 7.06 | -3.18 | -20.69 | -0.71 | 36.03 | -61.52 |
| 2 | -78.67 | -39.05 | 0.39 | -2.82 | -19.63 | -0.53 | 32.90 | -49.92 |
| 3 | -71.63 | -30.14 | 1.20 | -3.02 | -20.88 | -0.69 | 35.68 | -53.78 |
| 4 | -75.83 | -35.33 | 0.63 | -2.99 | -19.66 | -0.63 | 30.90 | -48.75 |
| 5 | -63.37 | -27.75 | 2.01 | -3.58 | -19.50 | -0.61 | 33.33 | -47.27 |
| 6 | -75.84 | -37.39 | 1.75 | -3.13 | -19.66 | -0.45 | 31.66 | -48.62 |
| 7 | -68.16 | -31.69 | 0.86 | -3.00 | -18.88 | -0.65 | 35.67 | -50.47 |
| 8 | -71.18 | -32.25 | -0.25 | -3.06 | -19.64 | -0.78 | 34.69 | -49.89 |
| 9 | -64.42 | -29.21 | 2.42 | -2.87 | -19.75 | -0.50 | 36.43 | -50.95 |
| 10 | -70.45 | -30.01 | 1.51 | -2.88 | -20.16 | -0.57 | 34.86 | -53.20 |
| 11 | -59.96 | -17.85 | 0.39 | -1.67 | -19.44 | -0.56 | 33.35 | -54.17 |
| 12 | -53.70 | -19.33 | 5.06 | -2.14 | -19.67 | -0.63 | 37.14 | -54.12 |
| 13 | -70.36 | -35.58 | 6.58 | -2.83 | -22.56 | -0.62 | 39.58 | -54.92 |
| 14 | -66.33 | -33.73 | 5.06 | -2.62 | -21.13 | -0.46 | 38.11 | -51.56 |
| 15 | -52.81 | -31.94 | 4.21 | -2.13 | -16.65 | -0.68 | 37.79 | -43.41 |
| 16 | -59.21 | -25.50 | 2.59 | -1.74 | -18.63 | -0.67 | 38.01 | -53.27 |
| 17 | -52.65 | -25.67 | 2.48 | -2.33 | -18.14 | -0.62 | 38.69 | -47.07 |
| 18 | -51.42 | -27.67 | 3.27 | -1.88 | -16.64 | -0.63 | 39.17 | -47.04 |
| 19 | -52.14 | -29.98 | 5.50 | -3.13 | -16.84 | -1.42 | 40.37 | -46.64 |
| 20 | -53.84 | -20.89 | 2.07 | -2.33 | -17.03 | -0.81 | 36.43 | -51.26 |
| 21 | -54.97 | -20.23 | 4.40 | -1.74 | -19.76 | -0.72 | 35.72 | -52.64 |
| 22 | -53.71 | -27.00 | 3.09 | -2.09 | -15.20 | -1.69 | 40.86 | -51.69 |
| 23 | -60.50 | -27.05 | 3.25 | -2.06 | -15.89 | -1.57 | 35.33 | -52.51 |
| 24 | -52.79 | -25.23 | 3.21 | -1.94 | -15.54 | -1.38 | 39.34 | -51.26 |
| 25 | -48.11 | -28.10 | 4.07 | -2.63 | -14.67 | -1.61 | 42.35 | -47.52 |
| 26 | -47.17 | -25.04 | 3.40 | -1.85 | -15.10 | -1.40 | 39.67 | -46.86 |
| 27 | -57.45 | -27.07 | 1.99 | -2.54 | -18.44 | -1.19 | 37.70 | -47.91 |
| 28 | -59.93 | -25.89 | 1.95 | -2.70 | -18.27 | -1.61 | 37.46 | -50.87 |
| 29 | -52.96 | -22.45 | 2.18 | -1.98 | -18.12 | -2.31 | 36.03 | -46.31 |
| 30 | -57.03 | -28.63 | 1.71 | -2.74 | -18.36 | -1.48 | 39.73 | -47.26 |
| 31 | -59.22 | -22.13 | 1.85 | -2.64 | -18.23 | -1.79 | 36.15 | -52.43 |
| 32 | -64.85 | -32.30 | 3.10 | -3.12 | -18.61 | -2.51 | 39.70 | -51.11 |
| 33 | -52.64 | -28.97 | 2.57 | -3.00 | -17.77 | -2.21 | 41.08 | -44.35 |
| 34 | -62.44 | -32.64 | 3.26 | -3.16 | -19.05 | -2.69 | 43.33 | -51.50 |
| 35 | -57.38 | -30.03 | 2.44 | -2.70 | -17.10 | -1.82 | 39.92 | -48.08 |
| 36 | -60.71 | -20.15 | 1.35 | -2.12 | -18.30 | -1.51 | 32.30 | -52.28 |
| 37 | -48.54 | -28.76 | 3.78 | -2.44 | -17.62 | -0.76 | 40.57 | -43.31 |
| 38 | -56.62 | -26.52 | 4.20 | -2.01 | -17.25 | -1.25 | 37.04 | -50.83 |
| 39 | -48.51 | -25.89 | 2.74 | -1.63 | -14.53 | -1.66 | 39.29 | -46.83 |
| 40 | -54.78 | -25.77 | 3.67 | -2.81 | -16.37 | -1.10 | 38.72 | -51.11 |
| 41 | -43.22 | -26.97 | 8.55 | -2.26 | -15.92 | -0.74 | 43.73 | -49.60 |
| 42 | -45.04 | -25.53 | 4.32 | -2.18 | -13.99 | -1.21 | 40.89 | -47.34 |
| 43 | -48.92 | -20.80 | 3.18 | -2.37 | -14.38 | -1.66 | 36.05 | -48.95 |
| 44 | -46.93 | -23.66 | 4.93 | -2.14 | -16.65 | -0.04 | 37.13 | -46.50 |
| 45 | -47.28 | -27.36 | 6.90 | -2.38 | -16.82 | -1.79 | 38.72 | -44.55 |
| 46 | -52.40 | -23.74 | 1.86 | -1.72 | -19.23 | -0.75 | 38.15 | -46.98 |
| 47 | -49.04 | -28.06 | 3.64 | -2.51 | -17.35 | -1.97 | 39.71 | -42.49 |
| 48 | -57.44 | -27.29 | 1.82 | -3.38 | -18.48 | -2.51 | 40.97 | -48.56 |
| 49 | -57.00 | -28.74 | 1.73 | -2.51 | -18.25 | -1.89 | 39.59 | -46.93 |
| 50 | -57.36 | -30.21 | 1.19 | -2.70 | -18.38 | -1.04 | 42.16 | -48.38 |
| 51 | -44.19 | -24.31 | 5.06 | -1.16 | -15.46 | -1.17 | 37.06 | -44.20 |
| 52 | -46.42 | -26.80 | 1.88 | -1.16 | -16.66 | -1.03 | 41.58 | -44.23 |
| 53 | -62.07 | -25.22 | 1.94 | -2.97 | -18.23 | -2.52 | 38.12 | -53.20 |
| 54 | -49.56 | -20.94 | 2.25 | -2.57 | -16.53 | -1.31 | 34.94 | -45.39 |
| 55 | -42.43 | -21.87 | 1.62 | -1.76 | -15.23 | -2.06 | 42.40 | -45.53 |
| 56 | -66.34 | -28.49 | 0.75 | -2.44 | -18.69 | -1.86 | 32.53 | -48.14 |
| 57 | -50.75 | -24.98 | 1.80 | -0.26 | -18.45 | -2.76 | 41.95 | -48.05 |
| 58 | -54.74 | -24.98 | 1.58 | -3.11 | -17.64 | -2.32 | 34.81 | -43.07 |
| 59 | -48.16 | -30.27 | 1.82 | -0.72 | -16.24 | -2.51 | 45.21 | -45.46 |
| 60 | -51.75 | -21.11 | 2.02 | -1.44 | -18.48 | -2.50 | 39.59 | -49.84 |
| 61 | -48.59 | -28.87 | 2.33 | -0.60 | -17.58 | -1.43 | 42.96 | -45.39 |
| 62 | -51.88 | -38.22 | 3.41 | -1.22 | -15.46 | -0.71 | 39.61 | -39.29 |
| 63 | -52.03 | -24.69 | 1.56 | -1.23 | -16.61 | -2.22 | 38.35 | -47.20 |
| 64 | -50.17 | -33.29 | 7.83 | -0.77 | -17.28 | -1.53 | 40.11 | -45.24 |
| 65 | -49.45 | -23.59 | 0.67 | -1.96 | -16.59 | -1.75 | 38.02 | -44.25 |
| 66 | -49.57 | -20.79 | 1.71 | -2.36 | -17.49 | -2.29 | 33.87 | -42.21 |
| 67 | -56.00 | -30.56 | 1.27 | -2.87 | -17.22 | -2.32 | 40.22 | -44.53 |
| 68 | -57.27 | -34.93 | 1.79 | -3.38 | -17.51 | -2.39 | 43.05 | -43.90 |
| 69 | -44.13 | -36.20 | 6.22 | -2.08 | -16.30 | -1.47 | 46.48 | -40.78 |
| 70 | -46.24 | -22.34 | 1.65 | -1.22 | -17.82 | -1.35 | 43.03 | -48.18 |
| 71 | -51.73 | -39.43 | 2.34 | -2.49 | -16.10 | -1.50 | 45.49 | -40.03 |
| 72 | -47.18 | -29.88 | 5.39 | -1.31 | -16.38 | -0.73 | 36.97 | -41.25 |
| 73 | -56.40 | -33.88 | 11.65 | -1.74 | -19.57 | -1.71 | 37.57 | -48.72 |
| 74 | -48.60 | -26.53 | 3.09 | -1.05 | -17.05 | -2.22 | 39.03 | -43.86 |
| 75 | -53.67 | -36.33 | 7.84 | -2.86 | -16.23 | -2.21 | 38.40 | -42.29 |
| 76 | -50.98 | -23.88 | 4.41 | -0.62 | -17.90 | -1.97 | 37.80 | -48.83 |
| 77 | -55.21 | -32.73 | 4.38 | -1.12 | -18.11 | -1.15 | 41.75 | -48.24 |
| 78 | -51.98 | -29.12 | 4.28 | -1.43 | -18.62 | -2.09 | 41.70 | -46.70 |
| 79 | -59.03 | -26.35 | 0.88 | -2.89 | -18.31 | -2.52 | 36.51 | -46.34 |
| 80 | -54.91 | -33.33 | 2.25 | -1.12 | -17.74 | -0.76 | 43.82 | -48.03 |
| 81 | -50.70 | -26.33 | 4.90 | -2.74 | -16.67 | -1.03 | 38.59 | -47.43 |
| 82 | -50.11 | -31.54 | 3.88 | -0.94 | -15.97 | -1.27 | 40.68 | -44.95 |
| 83 | -58.46 | -35.70 | 1.76 | -1.44 | -17.04 | -0.75 | 40.20 | -45.49 |
| 84 | -57.32 | -38.73 | 2.19 | -2.05 | -16.76 | -0.74 | 44.52 | -45.76 |
| 85 | -51.63 | -31.78 | 4.07 | -1.96 | -16.74 | -2.23 | 37.81 | -40.78 |
| 86 | -49.29 | -28.40 | 4.09 | -1.02 | -18.68 | -0.79 | 39.45 | -43.94 |
| 87 | -52.60 | -25.36 | 5.66 | -1.02 | -18.13 | -0.60 | 31.77 | -44.93 |
| 88 | -50.33 | -35.44 | 5.19 | -1.30 | -17.51 | -0.73 | 44.07 | -44.61 |
| 89 | -54.94 | -34.64 | 2.41 | -1.60 | -17.13 | -1.27 | 43.44 | -46.14 |
| 90 | -54.88 | -34.90 | 2.39 | -1.45 | -17.80 | -1.88 | 43.19 | -44.43 |
| 91 | -53.40 | -37.81 | 2.46 | -1.06 | -16.74 | -2.05 | 43.19 | -41.40 |
| 92 | -56.04 | -40.35 | 6.27 | -1.88 | -17.46 | -0.87 | 45.81 | -47.57 |
| 93 | -54.12 | -39.41 | 4.64 | -2.14 | -17.17 | -0.70 | 40.65 | -39.99 |
| 94 | -50.56 | -34.70 | 2.95 | -1.28 | -16.55 | -0.97 | 43.31 | -43.31 |
| 95 | -48.48 | -27.14 | 1.74 | -0.78 | -16.81 | -0.77 | 38.45 | -43.15 |
| 96 | -54.54 | -34.81 | 3.99 | -1.23 | -17.47 | -1.00 | 40.63 | -44.64 |
| 97 | -63.79 | -50.12 | 2.89 | -4.02 | -17.30 | -1.50 | 49.60 | -43.33 |
| 98 | -52.98 | -37.71 | 2.84 | -1.18 | -16.07 | -0.54 | 40.19 | -40.51 |
| 99 | -59.78 | -31.92 | 0.66 | -2.70 | -16.90 | -2.35 | 36.53 | -43.10 |
| 100 | -56.17 | -38.41 | 2.30 | -2.95 | -16.50 | -1.75 | 46.16 | -45.02 |
| AVG | **-55.25** | **-29.25** | **3.14** | **-2.12** | **-17.58** | **-1.37** | **39.11** | **-47.18** |
| MAX | **-78.67** | **-50.12** | **-0.25** | **-4.02** | **-22.56** | **-2.76** | **30.90** | **-61.52** |
| MIN | **-42.43** | **-17.85** | **11.65** | **-0.26** | **-13.99** | **-0.04** | **49.60** | **-39.29** |
| SD | **7.40** | **5.74** | **1.98** | **0.79** | **1.53** | **0.67** | **3.59** | **3.91** |

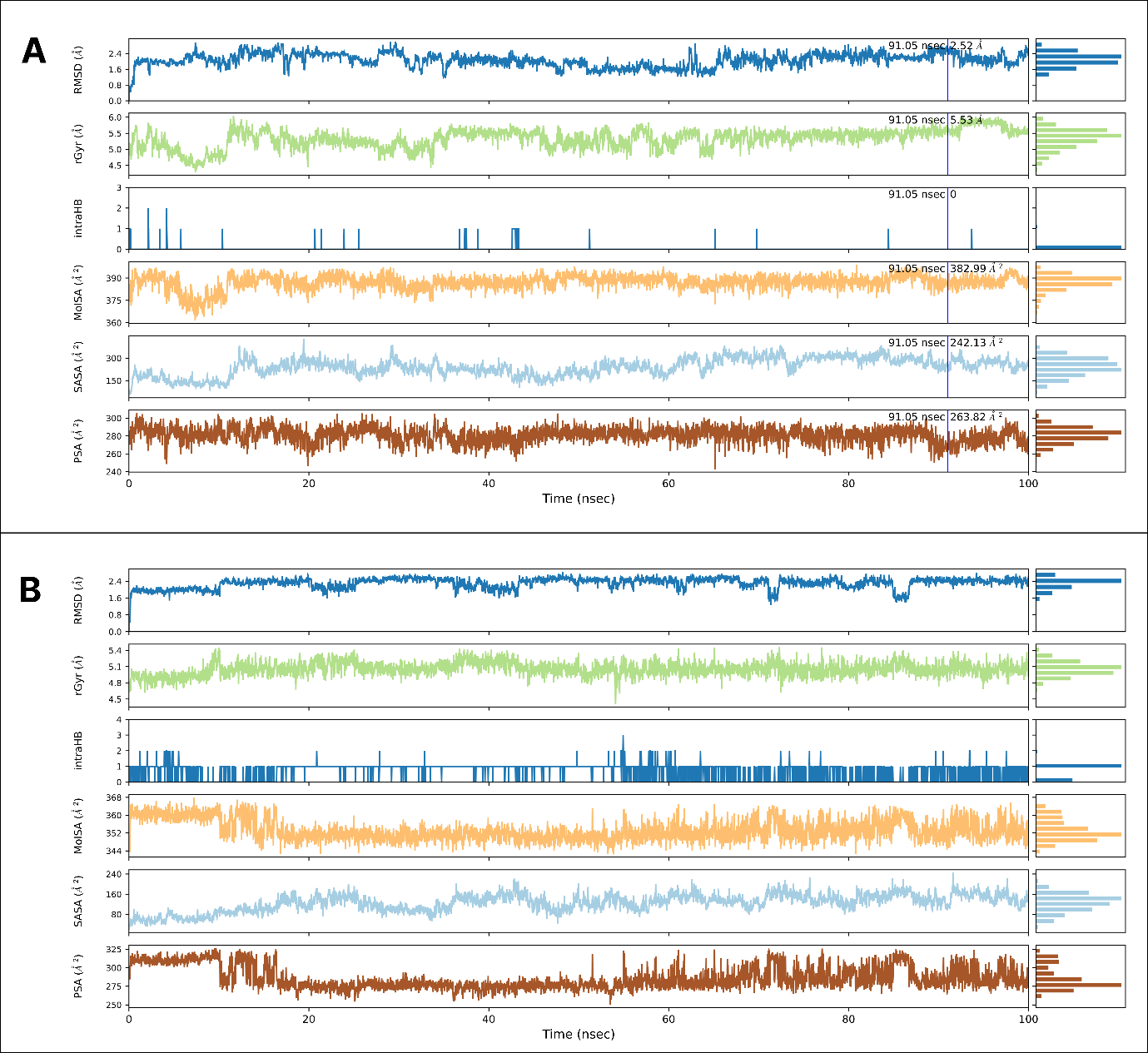

Figure 1 Variations in the properties RMSD, rGyr, MolSA, intraHB, SASA, and PSA for the two ligands (A: CNP0375130 and B: CNP0256178) during 100ns of MD simulation time.

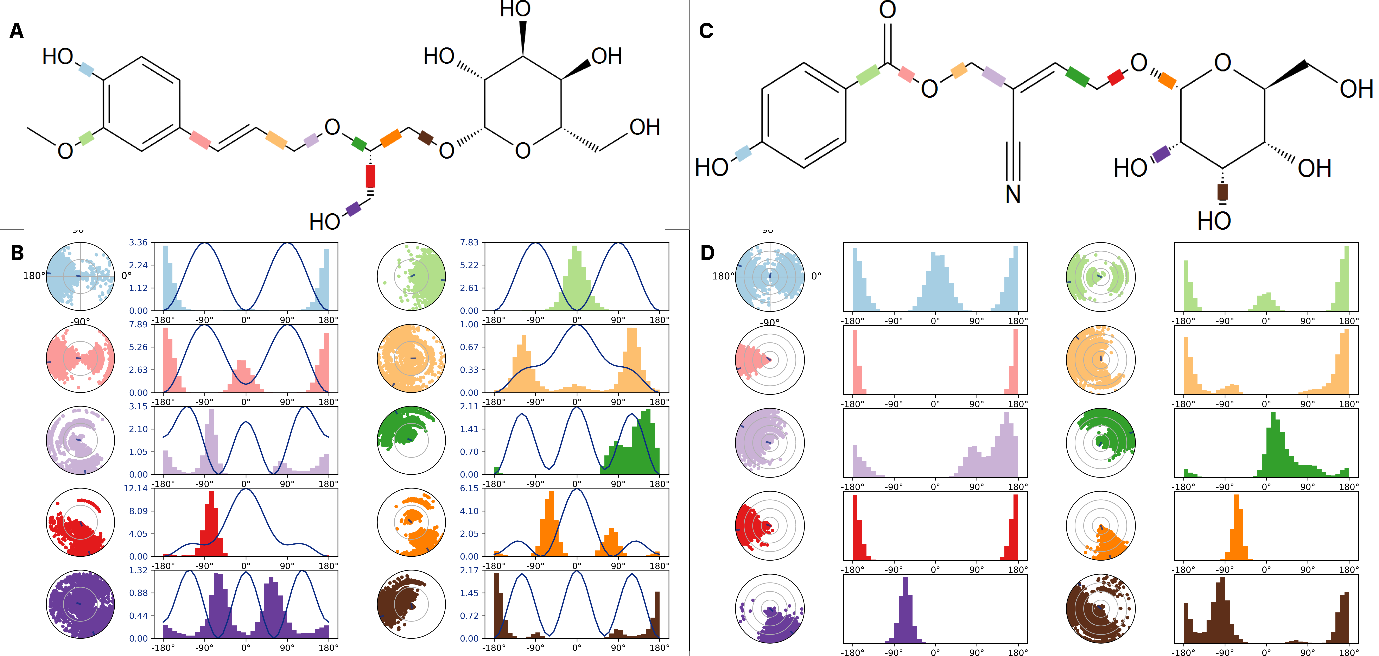

Figure 2 The ligand torsions plot summarizing the conformational evolution of every rotatable bond (RB) in the ligands. A: 2D schematic of CNP0375130 with color-coded rotatable bonds; B: Dial plot and bar plots of the rotatable bonds of ligand CNP0375130; C: 2D schematic of CNP0256178 with color-coded rotatable bonds; D: Dial plot and bar plots of the rotatable bonds of ligand CNP0256178

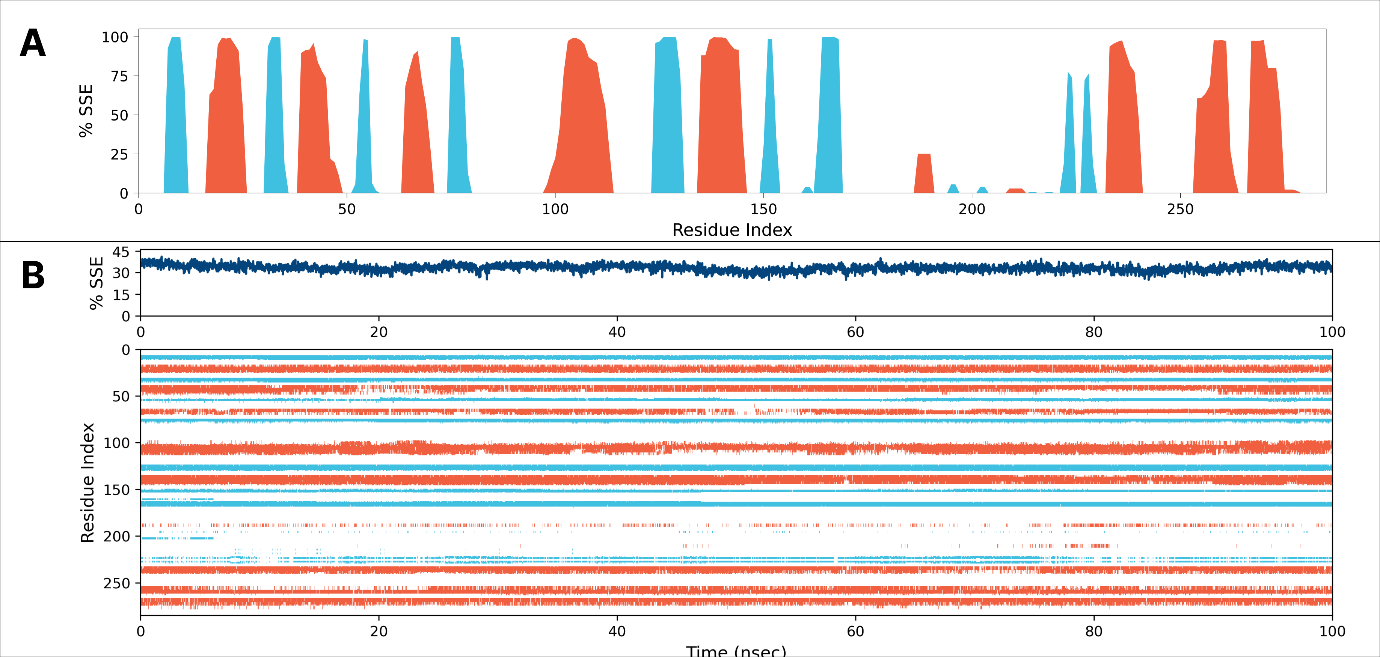

Figure 3 Protein secondary structure elements (SSE) including alpha-helices (in red) and beta-strands (in blue). A: SSE distribution by residue index throughout the protein structure. B: SSE composition for each trajectory frame throughout the simulation, and the plot at the bottom monitors each residue and its SSE assignment over time

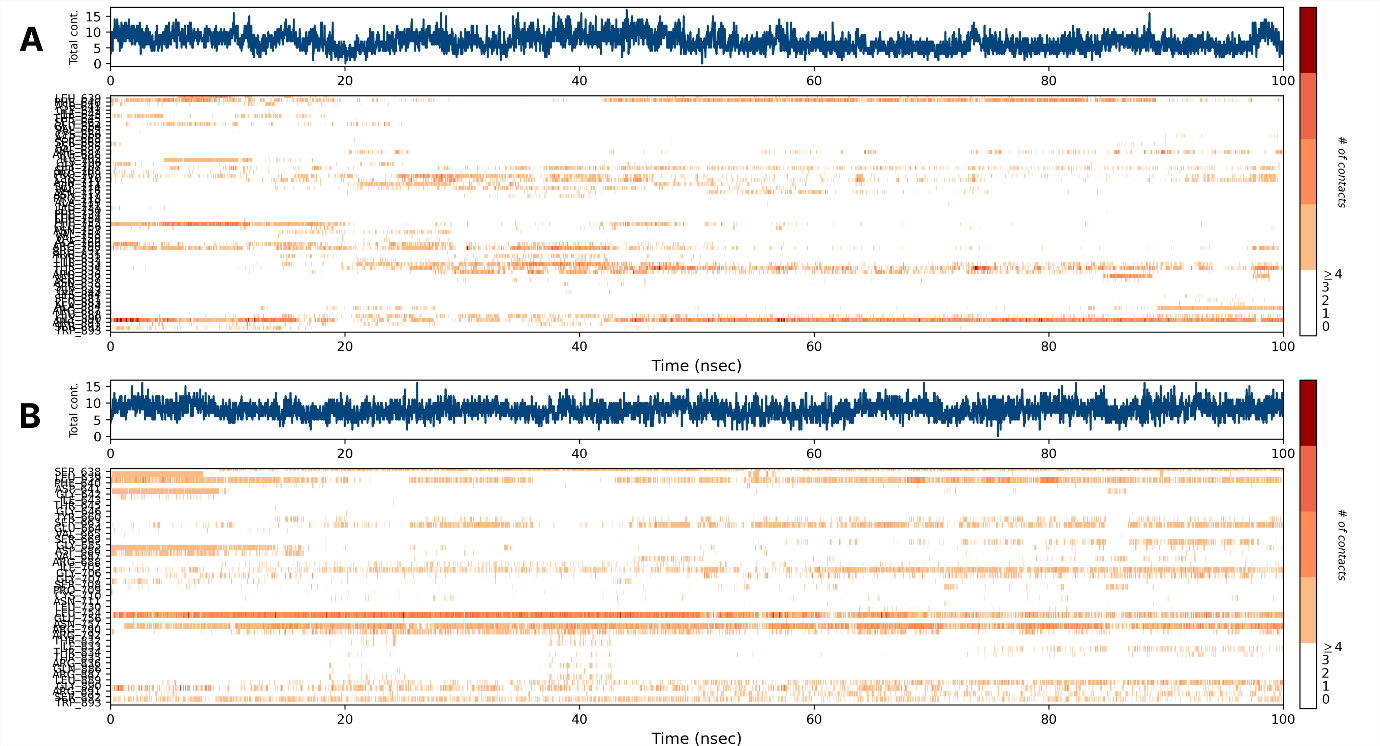

Figure 4 Timeline representations fraction of interactions between protein-ligand complex (A) 5YX2(A)-CNP0375130 (B) 5YX2(A)-CNP0256178. The top panel (in blue) shows the total number of specific contacts the protein makes with the ligand over the course of the trajectory. The bottom panel (in orange) shows which residues interact with the ligand in each trajectory frame.

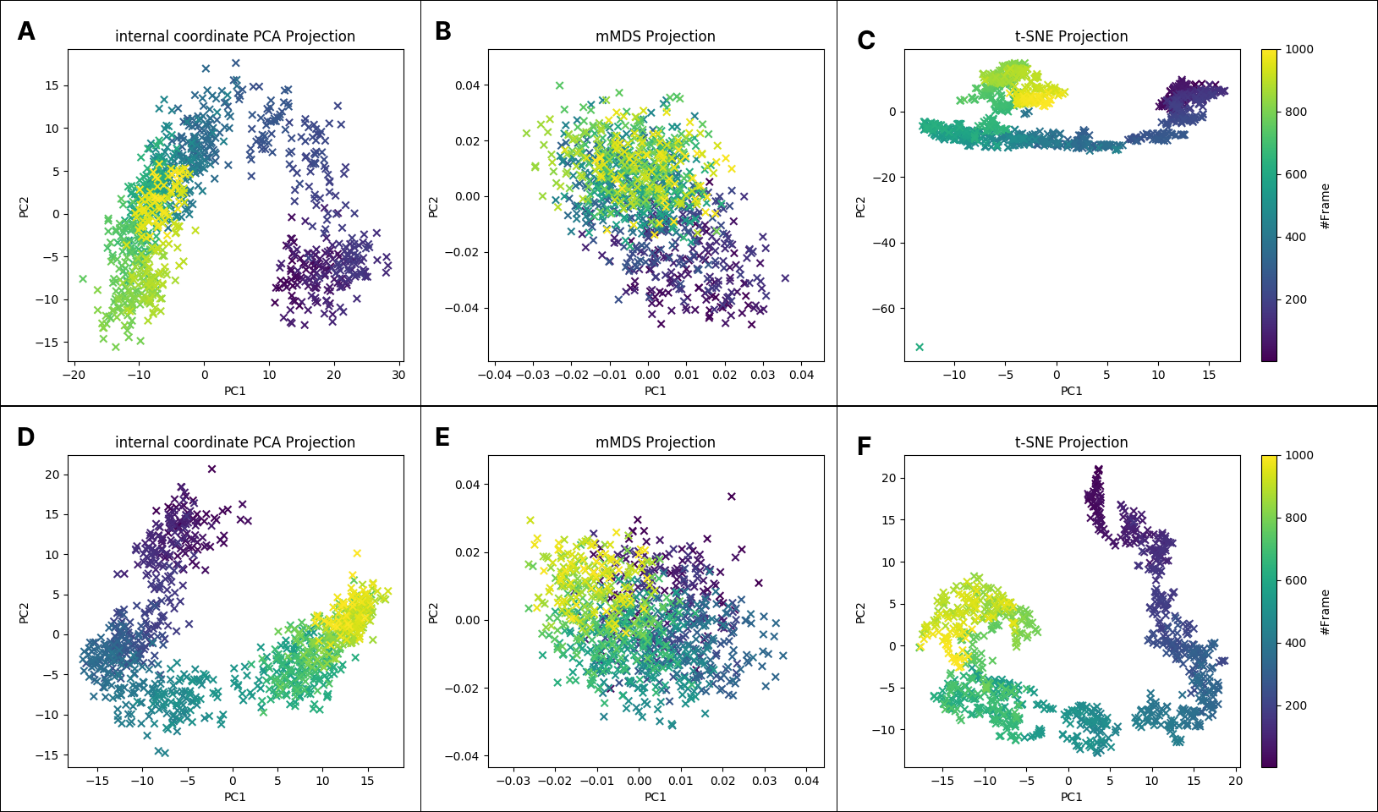

Figure 5 The Internal PCA, Multi-Dimensional Scaling (MDS) and t-distributed stochastic neighbour embedding (t-SNE) plots (A, B and C respectively for CNP0375130; D, E and F respectively for CNP0256178) illustrates principal components (PC2 vs. PC1) capturing variance in molecular dynamics data. Data points, colored by time, show conformational spread (wide vs. tight clusters), highlighting stable states (distinct clusters) and transitions (pathways). Density and color gradient indicate sampled conformations and temporal evolution.

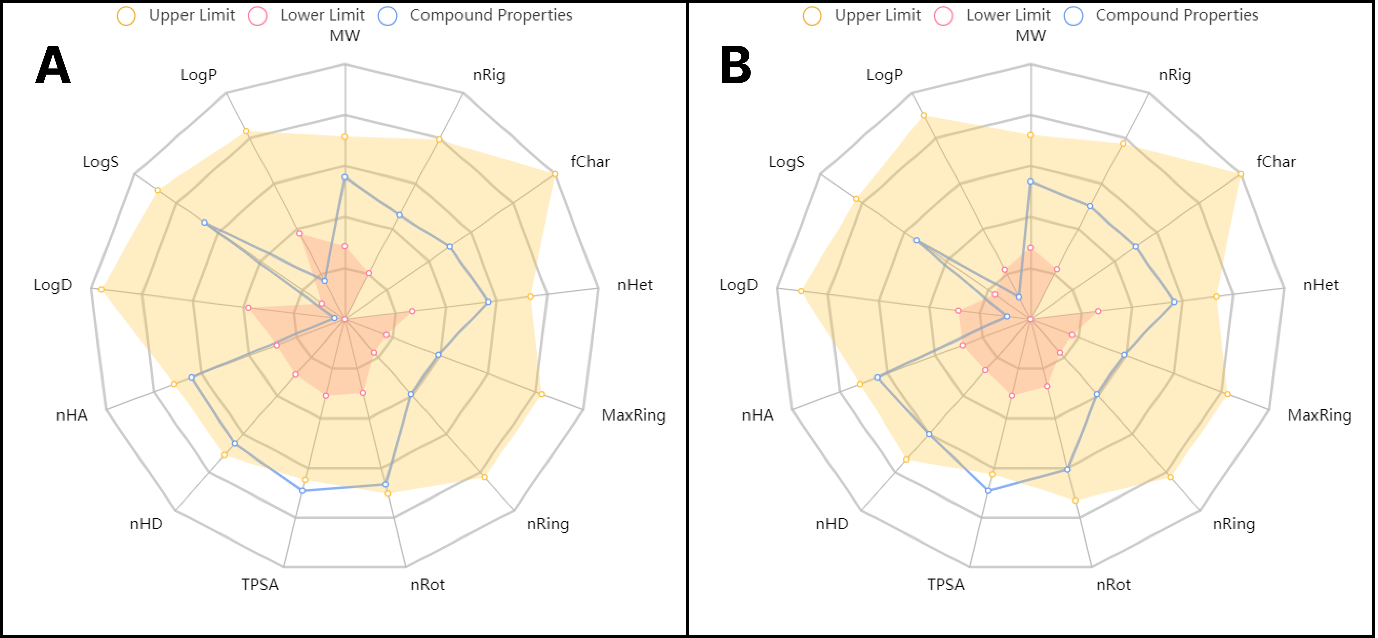

Figure 6 Bioavailability and Drug-likeliness of the two screened compounds generated from ADMETlab 2.0. A: CNP0375130; B: CNP0256178
